# Multiple phosphatidylinositol(3)phosphate roles in retinal pigment epithelium membrane recycling

**DOI:** 10.1101/2020.01.09.899815

**Authors:** Feng He, Melina A. Agosto, Ralph M. Nichols, Theodore G. Wensel

**Author notes:** Correspondence: Theodore G. Wensel, Verna and Marrs McLean Department of Biochemistry and Molecular Biology, Baylor College of Medicine, One Baylor Plaza, Houston, Texas 77030, USA;.

## Abstract

The low-abundance lipid phosphatidylinositol-3-phosphate (PI(3)P) is important for membrane dynamics in autophagy, endosome processing, and phagocytosis. In retinal pigmented epithelial cells (RPE) all three pathways are important, but phagocytosis of disk membranes shed from adjacent photoreceptors is especially important for ensuring health of photoreceptors and preventing retinal degeneration. By eliminating Vps34, the kinase responsible for synthesizing PI(3)P in RPE, we have found that PI(3)P plays distinct roles in each pathway. In phagocytosis it is not required for disk engulfment or phagosome transport but is essential for recruitment and lipidation of LC3. In contrast, initiation of autophagy and LC3 recruitment to autophagosomes does not require PI(3)P, which can be bypassed by an alternative mechanism of ATG16L recruitment that does not require PI(3)P, Rab11a, or WIPI2. In all three pathways, PI(3)P is essential for fusion with lysosomes; autophagosomes, phagosomes, and Rab7-positive late endosomes, as well as enlarged lysosomes, accumulate in large numbers in the absence of Vps34, leading to cell death.

## Introduction

In the retinal pigmented epithelium (RPE), two pathways for degradation of cellular material are essential for vision: autophagy, which engulfs and degrades intracellular components, and phagocytosis of photoreceptor outer segment (POS) tips (Young, 1967; Bok, 1993; Ferguson and Green, 2014). Both pathways have been studied sufficiently to reveal that these pathways use some of the same mechanisms and components so that they are to some extent antagonistic: one a response to a surfeit of material from outside the cell that must be digested, and the other a response to starvation and focused on homeostatic degradation of intracellular material. In addition to phagocytosis, RPE cells, like other cells, also capture nutrients and recycle membrane components through endocytosis. Much remains unknown about the details of all three pathways.

Phosphatidylinositol-3-phosphate (PI(3)P) has important roles in membrane trafficking pathways. It is synthesized by the class III PI-3-kinase, Vps34, which adds phosphate from ATP to the 3-position of the inositol ring in phosphtidylinositol (Raiborg et al., 2013). Recruitment of a complex containing Vps34 is an important early step in autophagy and phagocytosis, although complexes of different compositions are associated with each. In the canonical autophagy pathway, cellular stress activates the ULK1 complex, which in turn activates the PI3KC3 complex I, containing the scaffold protein Vps15, Atg14, Beclin 1 and the type III phosphoinositide 3-kinase, Vps34 (Stjepanovic et al, 2017); this activation leads to extension of a double-membrane phagophore enriched in PI(3)P (Axe et al, 2008; Hayashi-Nishino et al, 2009; Itakura and Mizushima, 2010; Kishi-Itakura et al, 2014). PI(3)P is thought to recruit WIPI2 (Polson et al, 2010; Dooley et al, 2014), followed by sequential recruitment of the two ligases, Atg7 (E1) and Atg3 (E2), and the complex of Atg16L1 with the Atg5-Atg 12 covalent conjugate (Yu, Chen, et al, 2018). Together these carry out a multistep ubiquitination-like process leading to covalent attachment of LC3 to phosphatidylethanolamine and formation of membrane-anchored LC3-II (Mizushima et al, 2001, 2003; Fujita et al, 2008; Lystad et al, 2019). Recent results suggest an even more complex picture, with robustness provided by redundant mechanisms such as starvation-induced LC3 lipidation in the absence of WIPI2 (Lystad et al, 2019) and PI(3)P (Ge et al., 2013). Both LC3-II and, separately, PI(3)P are thought to be important for subsequent steps of cargo attachment and ubiquitination, followed by transport to and fusion with lysosomes, where autophagosome contents are degraded. In Vps34 knockout cells, which are deficient in PI(3)P, LC3-II containing autophagosomes form, but do not colocalize with or fuse with lysosomes, and lysosomes proliferate in response (Zhou et al, 2010; Jaber et al, 2012; Bechtel et al, 2013; Reifler et al, 2014; He et al, 2016, 2019).

LC3 lipidation on phagosomes, referred to as LC3-associated phagocytosis (LAP), and the roles of Vps34 in it, are less well understood. In macrophages, LAP uses a subset of autophagy pathway components including LC3, ATG3, ATG7, ATG5, ATG12, and ATG16L1 (Martinez et al, 2011; Florey et al, 2011; Martinez et al, 2015; Fletcher et al, 2018; Sanjuan et al, 2007) as well as a Vps34-containing complex that includes Vps15, Beclin 1, UVRAG and RUBCN/Rubicon (RUN domain and cysteine-rich domain containing Beclin 1-interacting protein), but not Atg14 (Wong et al., 2018). LAP also diverges from autophagy in lack of involvement of the autophagy preinitiation complex, ATG13-ULK1-RB1CC1/FIP200-ATG101 (Kim et al, 2013; Muniz-Feliciano et al, 2017; Martinez et al, 2015).

RPE cells are professional phagocytes, ingesting and processing the distal 10% of the POS daily (Young, 1967; Bok, 1993). POS phagosome processing has been reported to involve LC3 recruitment and lipidation (Kim et al, 2013; Frost et al, 2015), with a specific requirement for the LC3B isoform (Dhingra et al., 2018), and to require the autophagy proteins ATG5, Beclin 1 (Kim et al. 2013) and RUBCN/Rubicon (Muniz-Feliciano et al, 2017). POS phagocyosis also activates mTORC1, a protein kinase complex which inhibits autophagy, as does Rubicon (Muniz Feliciano et al., 2017; Yu, Egbejimi, et al., 2018). The roles of Vps34 and PI(3)P in LAP of POS in RPE cells have not yet been explicitly explored, nor have their roles in upstream and downstream events such as engulfment, or transport to and fusion with lysosomes.

The roles of Vps34 and PI(3)P are even more poorly understood in endocytosis by RPE cells. PI(3)P was reported to play a critical role in actin polymerization at the surface of early endosomes in clathrin mediated endocytosis; its production was proposed to occur through action of a class II PI-3 kinase on PI(4)P followed by removal of the 4 phosphate (Daste et al., 2017). However, in those experiments endosome processing was blocked both by the non-selective PI-3 kinase inhibitor, wortmannin, and by a highly selective Vps34 inhibitor, suggesting that Vps34 is essential for this process as well. Consistent with this idea, we found previously that in rod cells and ON-bipolar cells of the retina, loss of Vps34 activity led to the accumulation of Rab7-labeled late endosomes (He et al., 2016, 2019).

To determine the roles of PI(3)P in membrane recycling pathways in the RPE, we combined *in vivo* studies in mice with experiments in immortalized RPE cell cultures. Tests of function were carried out *in vivo* using RPE-specific knockout of Vps34 and electroporation of fluorescent reporters along with immunofluorescence. These experiments were complemented by studies in cell culture using knockdown and reporter constructs as well as chemical treatments to dissect PI(3)P-dependent and -independent steps in each pathway. The results allow us to dissect common and distinct roles for Vps34 and PI(3)P in three different membrane recycling pathways in the same cells.

## Results

### Subcellular localization of PI(3)P and its depletion by knockout of Vps34 in RPE

Subretinal injection and electroporation of the PI(3)P-specific probe GFP-2xHrs (Gaullier et al, 1998; Gillooly et al, 2000; Furutani et al, 2006; He et al, 2016) in WT RPE resulted in labelling of many membrane structures ranging in size from small puncta to large vesicular structures in RPE cells *in vivo*, as expected from the known localization of PI(3)P to intracellular membrane compartments in other cell types (Fig 1A). Immunostaining with Rho antibody 1D4 revealed that a subset of these vesicles contain POS, confirming the presence of PI(3)P in phagosome membranes (Fig 1B,C). Interestingly, the transfected cells also exhibited an increased number of POS-containing phagosomes compared with neighboring untransfected cells (Fig 1B,C), suggesting that overexpression of the Hrs PI(3)P binding domains may sequester PI(3)P, competing with other PI(3)P-binding proteins, such as endogenous Hrs (Vieira et al, 2004), involved in phagosome maturation and slowing their removal by lysosomal degradation. In cultured RPE cells, PI(3)P colocalized primarily with a marker for the *trans*-Golgi, TGN46, and endosomal compartment markers Rab5 and Rab7, but not with markers for endoplasmic reticulum or the *cis*-Golgi (Fig S1A), as expected from the known localization of PI(3)P in other cell types (Gillooly 2000). Treatment with chloroquine to block autophagosome fusion with lysosomes led to accumulation of large numbers of autophagosomes marked by LC3 (Fig S1B), which were surrounded by large structures enriched in PI(3)P, as found in many other cell types.

**Figure 1.**
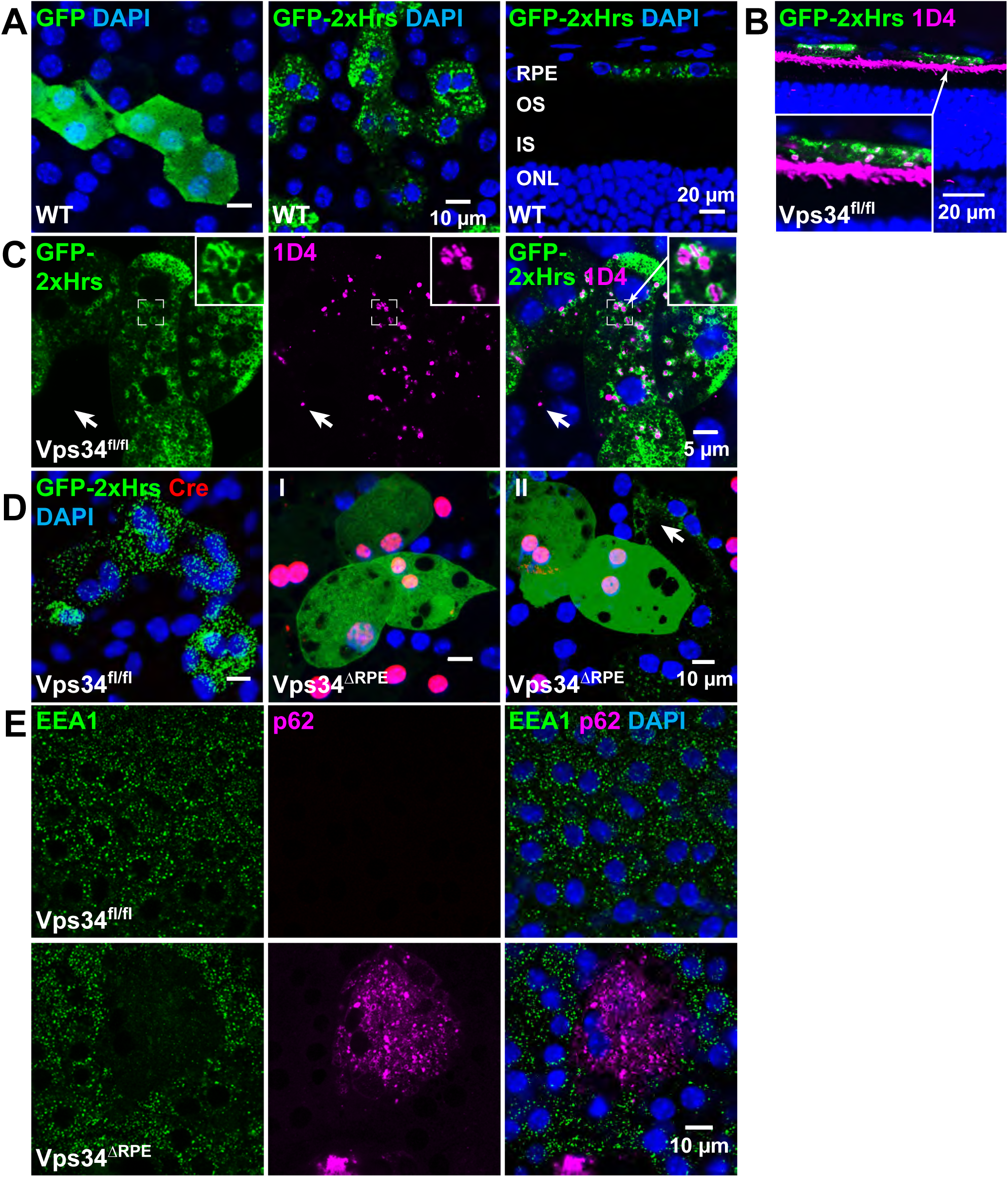
PI(3)P subcellular localization in RPE and depletion in Vps34 KO. (**A**) Plasmids expressing GFP (A, far left panel) or the PI(3)P probe GFP-2xHrs (A-D) were injected subretinally and electroporated to transfect RPE cells. Flatmount RPE (A, middle panel) and vertical eyecup sections (A, far right panel) confirmed expression in the RPE layer and punctate localization of GFP-2xHrs. (**B, C**) Eyecup section (B) and RPE flatmount (C) from control mice injected with GFP-2xHrs to mark PI(3)P and labeled with 1D4 antibody to mark POS phagosomes in RPE. (**D**) Cre immunostaining in flatmount electroporated RPE revealed mosaic expression of Cre in the Vps34^ΔRPE^ mice. Punctate localization of GFP-2xHrs was lost in Cre-positive KO cells (*i, ii*), but maintained in Cre-negative cells (arrow). The green channel is over-saturated in *ii* to show the dimmer puncta in Cre-negative cells. (**E**) Punctate endosomal localization of PI(3)P-binding protein EEA1 was disrupted in KO cells, identified by accumulation of p62

To knockout Vps34 and test the function of PI(3)P in RPE, we crossed mice carrying a Cre transgene under control of the promoter for bestrophin 1 (*BEST1*) (Iacovelli et al, 2011) with a strain containing a “floxed” allele of the mouse *Pik3c3* gene encoding Vps34, with loxP sites flanking exons 17-18, resulting in an in-frame deletion of the Vps34 ATP-binding domain upon expression of Cre (Zhou 2010) in the post-natal RPE. Cre immunostaining and probing for PI(3)P confirmed expression and Vps34 knockout in an apparently random subset of RPE cells (Vps34^ΔRPE^), consistent with the reported patchy mosaic expression of Cre from this transgene (Fig 1C). In Cre-positive (i.e. KO) cells the GFP-2xHrs probe was diffusely localized, indicating loss of the Vps34 product, PI(3)P (Fig 1C). Similarly, punctate immunostaining of the endogenous PI(3)P binding protein EEA1 in endosomes was greatly reduced in KO RPE (Fig 1D), and KO cells accumulated many puncta positive for the autophagy protein p62 as observed previously in Vps34 KO neurons (Zhou et al, 2010; He et al, 2016, 2019). These results confirm both the specificity of the PI(3)P probe and the success of the knockout in Cre-expressing cells.

### Loss of PI(3)P leads to accumulation of abnormal membrane structures and RPE cell degeneration

Consistent with the mosaic expression of Cre, a subset of RPE cells in Vps34^ΔRPE^ mice had altered morphology (Fig. 2). Patches of RPE lacking tight junctions marked by ZO-1 and with reduced number of nuclei indicate severe cellular pathology and death in Vps34 knockout cells (Fig 2A). Transmission electron microscopy (TEM) of retinal sections revealed an abnormal RPE layer with cells enlarged to varying degrees and containing accumulated vesicles of abnormal structure, including multivesicular structures (Fig 2B, S2). Secondary effects on photoreceptor health are expected as a result of RPE degeneration, and accordingly, we observed severely disordered photoreceptor outer segments (POS) in retinas from Vps34^ΔRPE^ mice (Fig 2B). We did not observe significant differences in average outer nuclear layer (ONL) thickness, but cannot rule out some photoreceptor cell death in regions adjacent to the most severely degenerated RPE. These observations confirm that in the RPE Vps34 and PI(3)P are essential for normal membrane structure and cell health.

**Figure 2.**
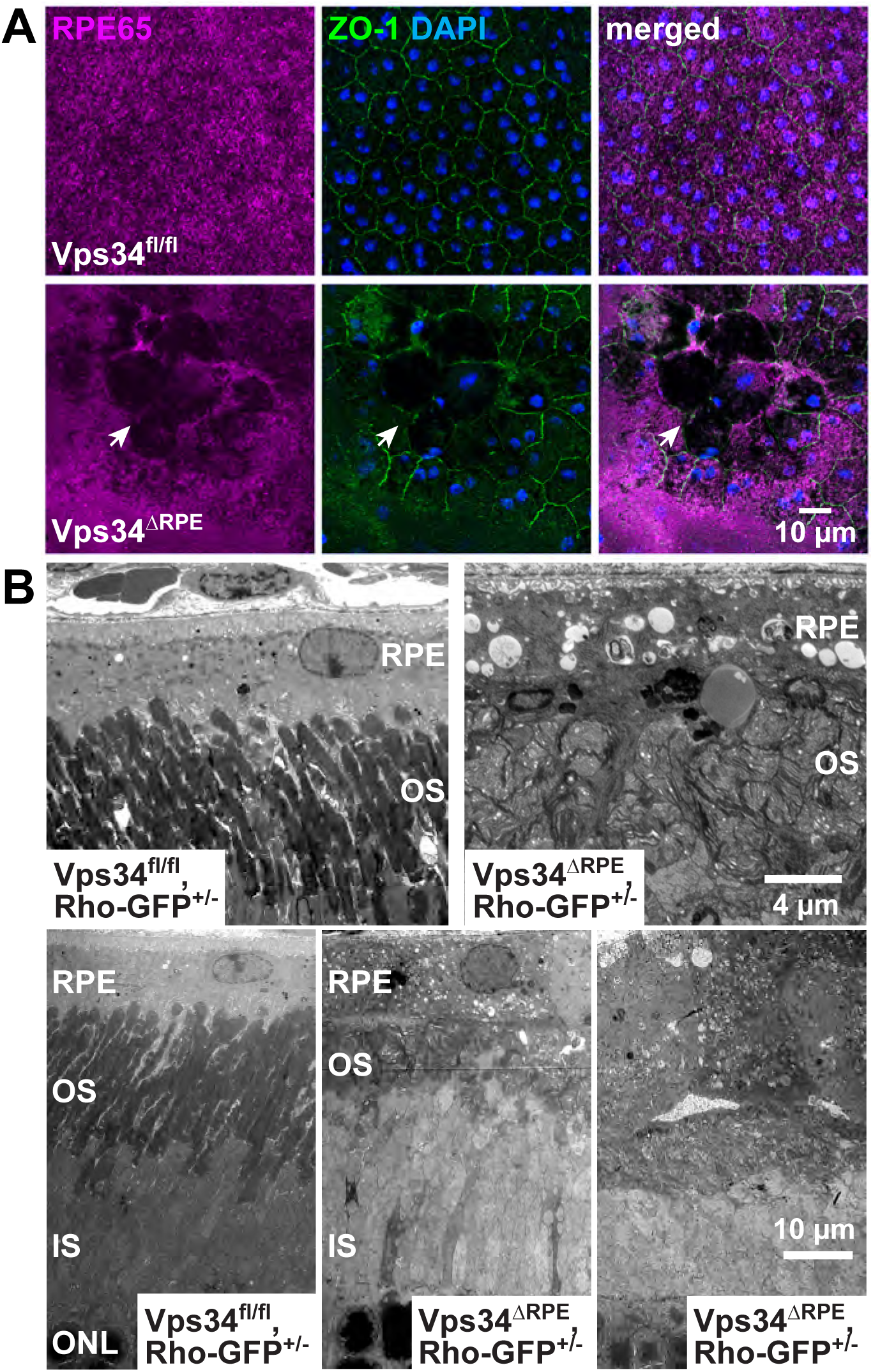
Vps34^ΔRPE^ mice exhibit RPE cell degeneration. (**A**) Immunostaining for RPE65 and the tight junction protein ZO-1 in RPE flatmounts indicate abnormal morphology and cell death in the KO mice. (**B**) Abnormal morphology of both RPE cells and photoreceptor outer segments was observed by TEM of eyecup sections.

### POS phagosomes form but do not recruit LC3 in the absence of PI(3)P

One of the vital roles of the RPE is daily phagocytosis and degradation of POS tips (Young, 1967; Bok, 1993). To examine phagosomes, eyes were harvested within 2 hours of light onset in the morning, during the peak of POS phagocytosis (Goldman et al, 1980).

To aid in visualizing phagosomes, we produced Vps34^ΔRPE^ or control Vps34^fl/fl^ mice heterozygous for a low-expressing allele of human Rho-GFP (Chan et al, 2004). Small numbers of fluorescent phagosomes could be observed in the control mice, depending on time of day, but regardless of what time the eyes were collected, accumulation of Rho-containing puncta was observed in a subset of RPE cells in the Vps34^ΔRPE^ KO mice coinciding with Cre expression (Fig 3A,B). This result indicates that Vps34 and PI(3)P are not required for engulfment of POS, but that there is a block to a later step needed for phagosome clearance. TEM confirmed the accumulation of compartments containing POS disk membranes (Fig 3C, S2). In control mice, the small number of Rho-containing phagosomes observed in RPE cells colocalized with LC3, indicating the presence of lipidated LC3 (Fig 4A), as reported previously for POS phagocytosis (Kim et al, 2013; Frost et al, 2015) as well as for LC3-associated phagocytosis in other cell types (Heckmann and Green, 2019; Martinez et al, 2015). In contrast, in the KO RPE, LC3 did not colocalize with the accumulated Rho puncta, revealing defective LC3 recruitment and, presumably, defective lipidation on POS-containing phagosomes, in the absence of PI(3)P (Fig 4A). Consistent with these results, LC3 colocalized with Rho puncta in cultured ARPE-19 cells fed with purified Rho-GFP-containing POS, and treatment with the Vps34 selective inhibitor Vps34-IN1 (Bago et al, 2014) suppressed LC3 recruitment (Fig 4C). The KO RPE contained a greatly increased number of large LAMP2-containing membranes, indicating upregulation of lysosomes, as observed in other cell types lacking Vps34. Strikingly, even with the large numbers of both phagosomes and lysosomes, there was virtually no colocalization of the two, indicating deficient phagosome trafficking and lysosome fusion in the absence of PI(3)P (Fig 4D).

**Figure 3.**
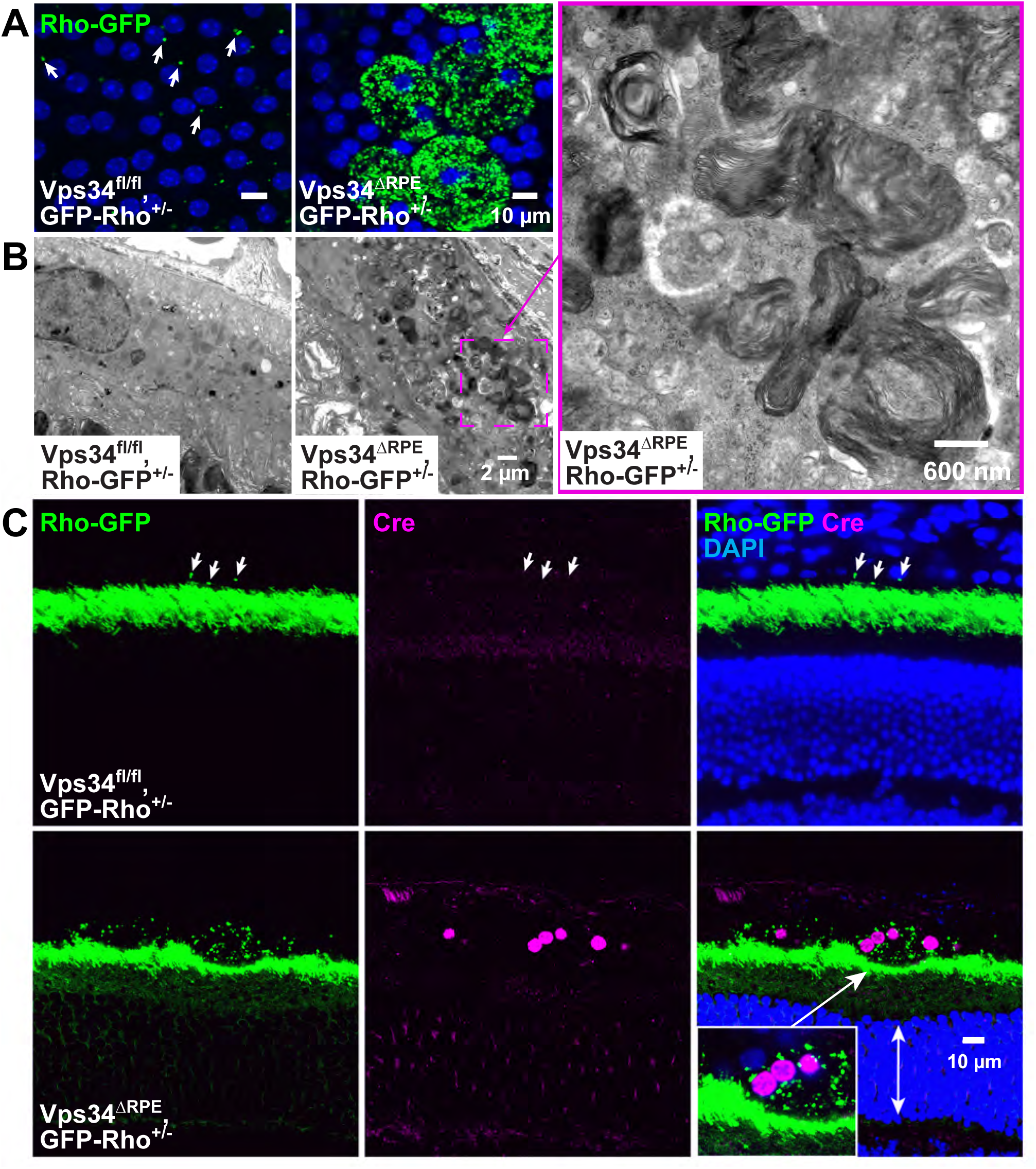
Phagosome processing is impaired following POS engulfment in Vps34 KO RPE. (**A**) RPE flatmount from control (left) and Vps34^ΔRPE^ mice (right) also expressing Rho-GFP show accumulation of Rho-containing phagosomes in KO RPE cells. (**B**) TEM from a region containing accumulated POS-containing phagosomes. (**C**) Eyecup sections from mice of same genotypes as in A, with immunostaining for Cre, showing the highest number of phagosomes in the Cre-positive regions.

**Figure 4.**
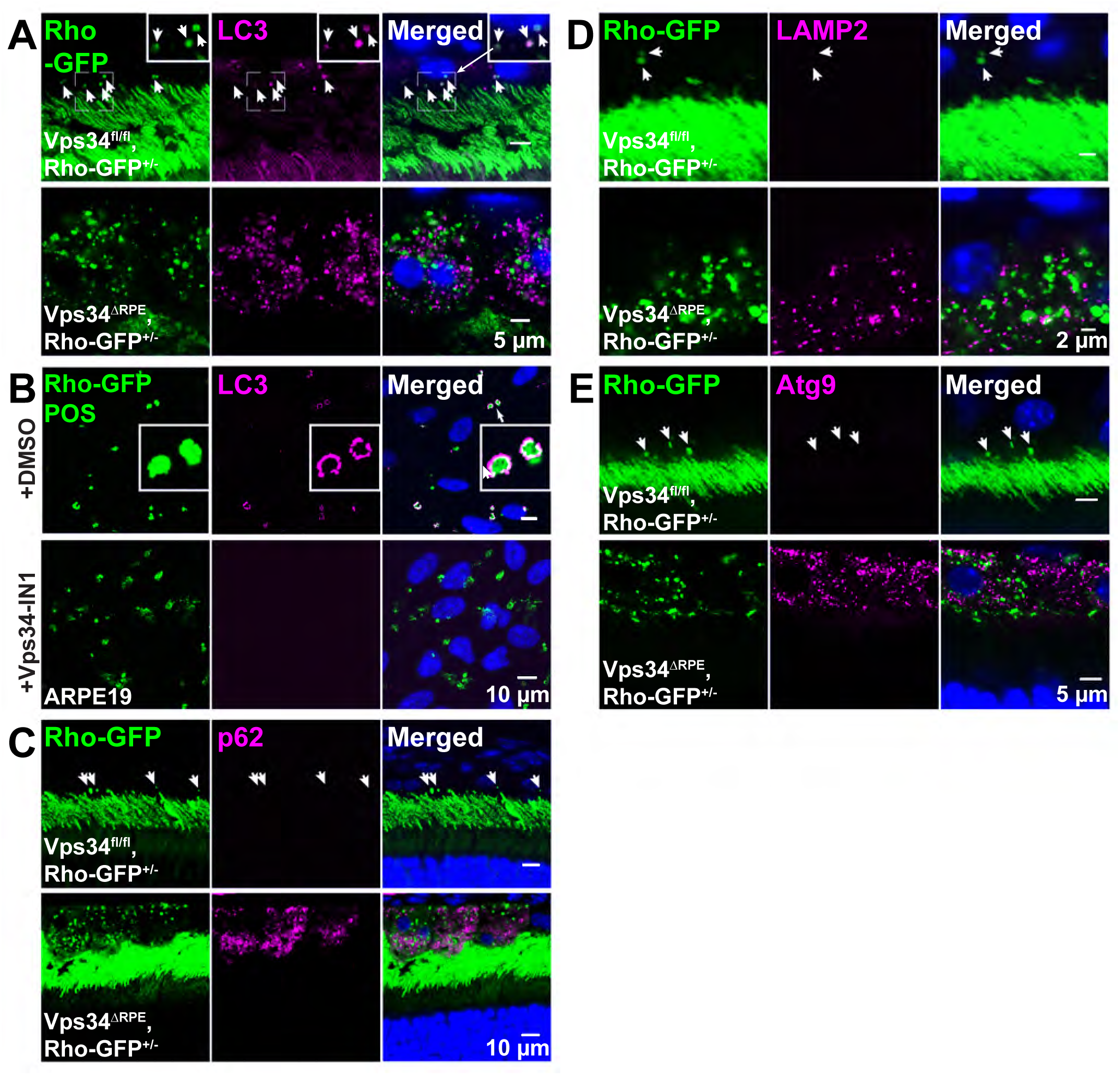
Phagosome maturation in Vps34 KO RPE is impaired prior to LC3 lipidation. (**A**) In eyecup sections (A) immunostained with LC3 antibody, Rho-containing phagosomes colocalized with LC3 in control, but not KO, RPE. (**B**) *In vitro* phagocytosis assay with ARPE-19 cells. LC3 colocalized with fed OS; Vps34-IN1 treatment suppressed LC3 recruitment. (**C-E**) Phagosomes did not colocalize with p62 (C), lysosome marker, LAMP2 (D) or Atg9 (E) in control or KO RPE.

LAP in general, and POS phagocytosis in RPE cells in particular, involves a subset of components of the autophagy pathway and has been referred to as “non-canonical autophagy” (Kim et al, 2013). In RPE, these include Atg5 and Beclin 1, but not Atg13, FIP200 or ULK1 (Kim et al, 2013). We tested POS phagosomes in RPE for the presence of two autophagy markers, the transmembrane autophagy protein Atg9 and the LC3-II-binding cargo adaptor, p62 (Fig 4D,E). We could not detect their presence in phagosomes in either WT or Vps34 KO RPE, although both accumulated on autophagosomes in the knockout cells (see below). Thus, beyond the lipidation of LC3, which differs in its requirement for PI(3)P from autophagy, phagosome processing in RPE appears to follow a distinct pathway from autophagy prior to lysosome fusion.

Rho in RPE phagosomes has been reported to undergo sequential degradative processing during phagosome maturation – first proteolytic degradation of the cytoplasmic C-terminus, associated with an endosome interaction, followed by loss of an intradiscal N-terminal epitope upon lysosome fusion (Wavre-Shapton et al, 2014; Esteve-Rudd et al, 2014). To probe for selective degradation, KO and control sections were co-immunostained with Rho mAbs 4D2 and 1D4, which recognize N- and C-terminal epitopes, respectively and can be distinguished by isotype-specific secondaries. In both genotypes, examples were observed of Rho-containing phagosomes labeled with both antibodies, preferentially with 4D2, or preferentially with 1D4, suggesting heterogeneous post-engulfment compartments differing in relative degradation rates or epitope accessibility (Fig S3).

### LC3-positive autophagosomes accumulate in knockout RPE

Vps34 function is required for completion of autophagy, and knockout of Vps34 has been shown to result in arrest prior to autophagolysosome formation in every cell type tested (Zhou et al, 2010; Jaber et al, 2012; Bechtel et al, 2013; He et al, 2016, 2019; Grieco et al., 2018). Similarly, in the KO RPE, puncta containing LC3 and autophagy proteins Atg9 and p62 accumulated (Fig 4A,E,F, 5A,E) and colocalized (Fig 5C,D), along with ubiquitinated proteins (Fig 5G). In TEM images of KO RPE, regions with numerous membrane structures were observed including double-membrane and multilamellar vesicles (Fig 5B,S2), consistent with autophagosomes stalled prior to lysosome fusion.

**Figure 5.**
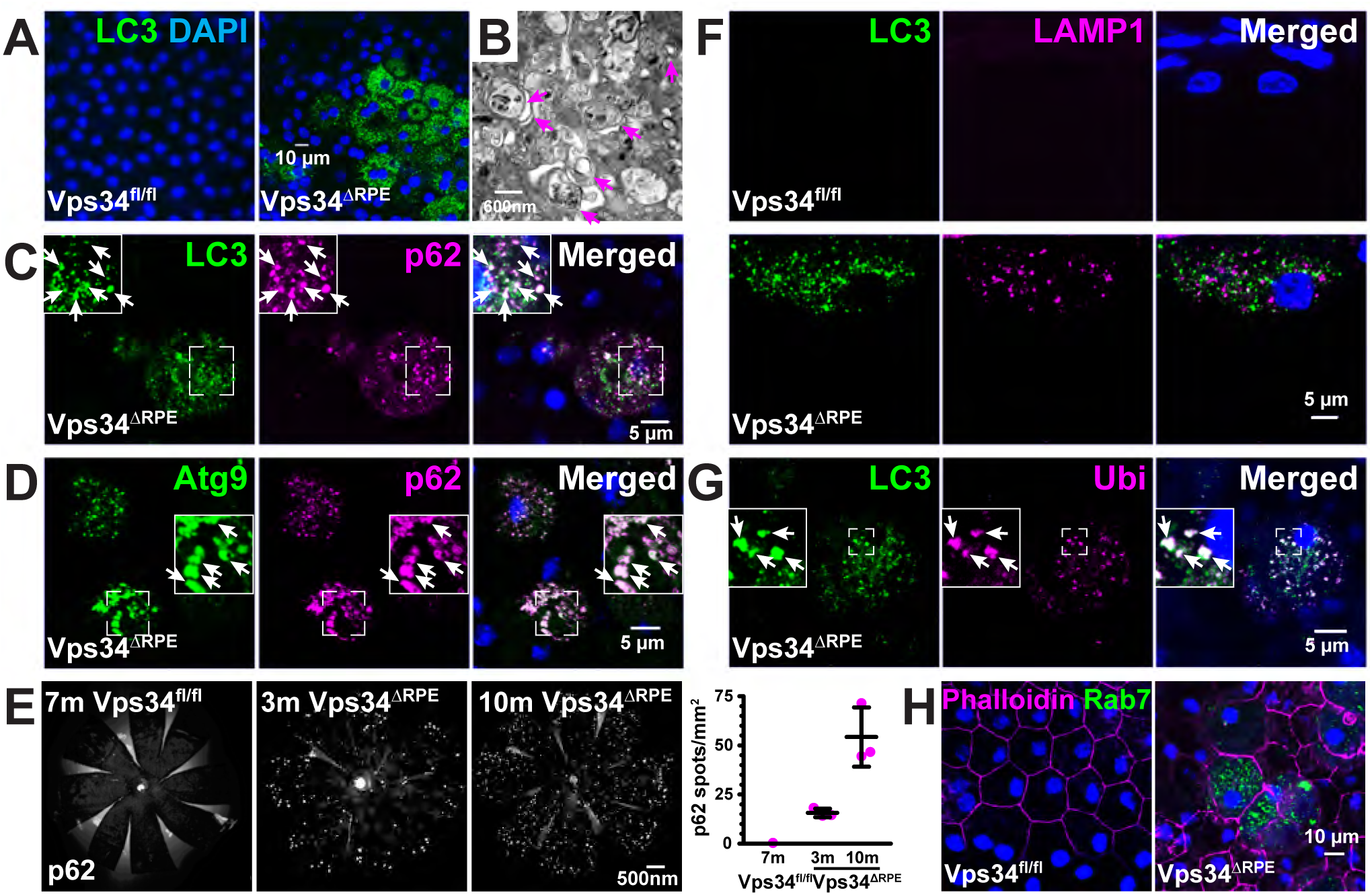
Autophagosome maturation in Vps34 KO RPE is impaired post-LC3 lipidation. (**A**) LC3-positive puncta accumulated in KO RPE. (**B**) TEM from region containing accumulated multi-membraned vesicles (arrows). (**C,D,G**) LC3 puncta accumulated and colocalized with p62 (C) and ubiquitin (G), and p62 colocalized with Atg9 (D). (**F**) LC3 and LAMP1 puncta did not colocalize, indicating impairment prior to autophagosome-lysosome fusion. (**E**) (left) p62-positive puncta accumulated in an age-dependent manner in KO RPE. (right) Quantification of images as shown in (E) from 3m and 10m old Vps34^ΔRPE^ mice. (**H**) Rab7 puncta accumulated in KO RPE.

In contrast to phagosomes, in which LC3 was absent in the KO (Fig 4A), the autophagosome markers colocalized with the accumulated LC3 puncta (Fig 5C,D,G). As indicated above there was a proliferation of enlarged lysosomes, but these did not co-localize with the LC3-positive autophagosomes (Fig 5F), consistent with a defect in the autophagy pathway after LC3 recruitment and lipidation but prior to autophagosome-lysosome fusion. Our data are consistent with reports of normal or increased levels of LC3 recruitment to autophagosomes and of lipidated LC3 in the absence of Vps34 function in other cell types (Zhou et al, 2010; Jaber et al, 2012; Bechtel et al, 2013; Reifler et al, 2014; He et al, 2016; Lystad et al, 2019),

### WIPI2 and PI(3)P are dispensable for LC3 recruitment to autophagosomes

LC3 recruitment is thought to occur via sequential recruitment of 1) WIPI2 via binding to PI(3)P; 2) Atg16L1 binding to WIPI2; and 3) Atg5-12, Atg3, and LC3 (Itakura and Mizushima, 2010; Dooley et al, 2014). To examine the mechanism of PI(3)P-independent LC3 recruitment, we used cultured human RPE cell lines hTERT-RPE-1 (hereafter hRPE1) and ARPE-19 as models.

In ARPE-19 cells in normal growth media, a small of number of WIPI2 puncta, and no LC3 puncta, were observed (Fig 6A). Overnight CQ treatment resulted in large increases in the number of both LC3 puncta and WIPI2 puncta, as expected, but they did not colocalize (Fig 6B), suggesting that after WIPI2 has recruited the complex responsible for conjugation of LC3-I to phosphatidylethanolamine, it and LC3-II segregate to different membrane compartments. Overnight treatment with the Vps34 inhibitor Vps34-IN1 (Bago et al, 2014) abolished the basal WIPI2 puncta seen without CQ, as well as the accumulated WIPI2 puncta in the presence of CQ (Fig 6C,D). These data show that PI(3)P is required for WIPI2 recruitment, consistent with previous reports (Dooley et al, 2014; Polson et al, 2010; Karanasios et al, 2013; Brier et al, 2019). LC3 puncta, in contrast, were still present in Vps34-IN1-treated cells (Fig 6D), consistent with a PI(3)P- and WIPI2-independent route for LC3 recruitment. Similar results were seen with overnight CQ and Vps34-IN1 treatment of DsRed-WIPI2-transfected hRPE1 cells (Fig S4C,D). However, in cells treated overnight with CQ, then treated acutely with Vps34-IN1 for 90 min, DsRed-WIPI2 puncta remained, whereas GFP-2xHrs puncta disappeared, verifying PI(3)P depletion (Fig S4C,D). Similarly, live cell imaging of CQ-treated hRPE1 cells transfected with GFP-2xHrs or GFP-WIPI2 showed that upon addition of Vps34-IN1, GFP-2xHrs rapidly dissociated from intracellular compartments as expected (Fig S4A), verifying depletion of PI(3)P from the membranes. In contrast, GFP-WIPI2 remained stably associated for at least 90 minutes, despite PI(3)P depletion (Fig S4B). Taken together, the observations that overnight treatment with Vps34-IN1 abolished WIPI2 puncta formation, while acute treatment failed to elute WIPI2 from punctate structures, suggest that PI(3)P is required for WIPI2 recruitment to autophagosome membranes, but that once it is bound, other interactions are sufficient to maintain its localization.

**Figure 6.**
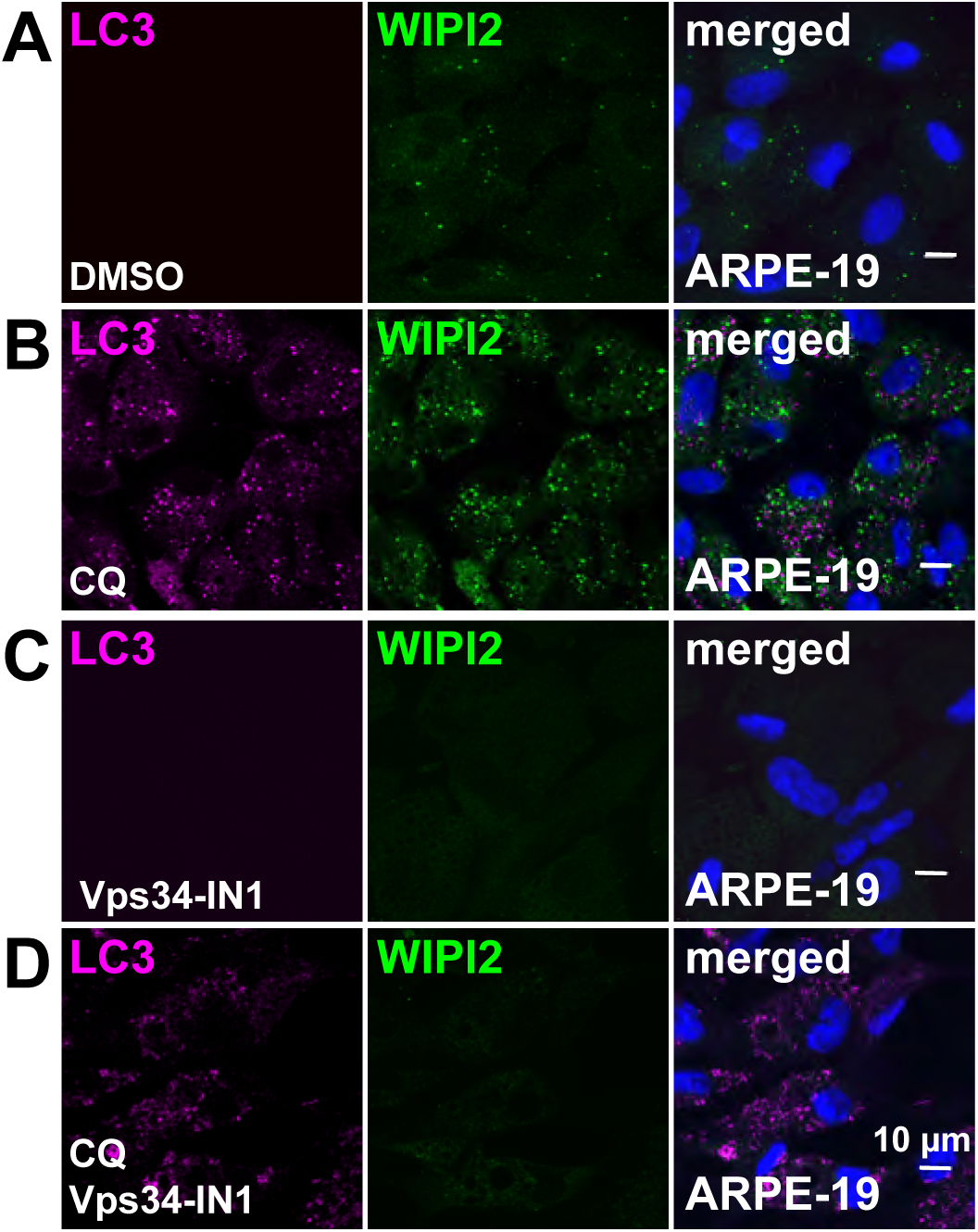
WIPI2 puncta formation requires PI(3)P, but LC3 puncta formation does not. (**A-D**) ARPE-19 cells were treated overnight with vehicle control (DMSO), Vps34-IN1, CQ, or both Vps34-IN1 and CQ together, then immunostained for LC3 and WIPI2.

WIPI2 is a member of the PROPPIN family, which is well known for binding to both PI(3)P and PI(3,5)P2 (Dove et al, 2004, 2009; Liang et al, 2019), and human WIPI2 was reported to bind PI(3)P, PI(3,5)P2, PI(4)P, and PI(5)P in lipid overlay assays (Liang et al, 2019; Vicinanza et al, 2015). However, we found that mouse GST-WIPI2 phosphoinositide binding was highly specific for PI(3)P, although binding was weak compared to GST-2xHrs (Fig 7A). In titrations of PI(3)P in PI-ELISA assays (He et al, 2016) GST-WIPI2 had a ∼6-fold lower affinity for PI(3)P than GST-2xHrs (*K_1/2_* values were 2.2 x 10^-3^ (±1.8 x10^-4^) and 3.8 x 10^-4^ (±3.9 x 10^-5^)(SEM, n=3) mol PI(3)P/mol total phospholipid, for WIPI2 and 2xHrs, respectively; corresponding to surface densities of roughly 494 or 2800 molecules/µm^2^, assuming a nominal area per phospholipid of 77 A^2^ (Burke et al., 1973) (Fig 7C). WIPI2 has two PI binding sites coordinated by the conserved FRRG motif (Baskaran et al, 2012; Krick et al, 2012; Liang et al, 2019). Mutation of one (WIPI2^FTRG)^ or both (WIPI2^FTTG^) of the PI binding sites virtually eliminated binding to PI(3)P (Fig 7A,B).

**Figure 7.**
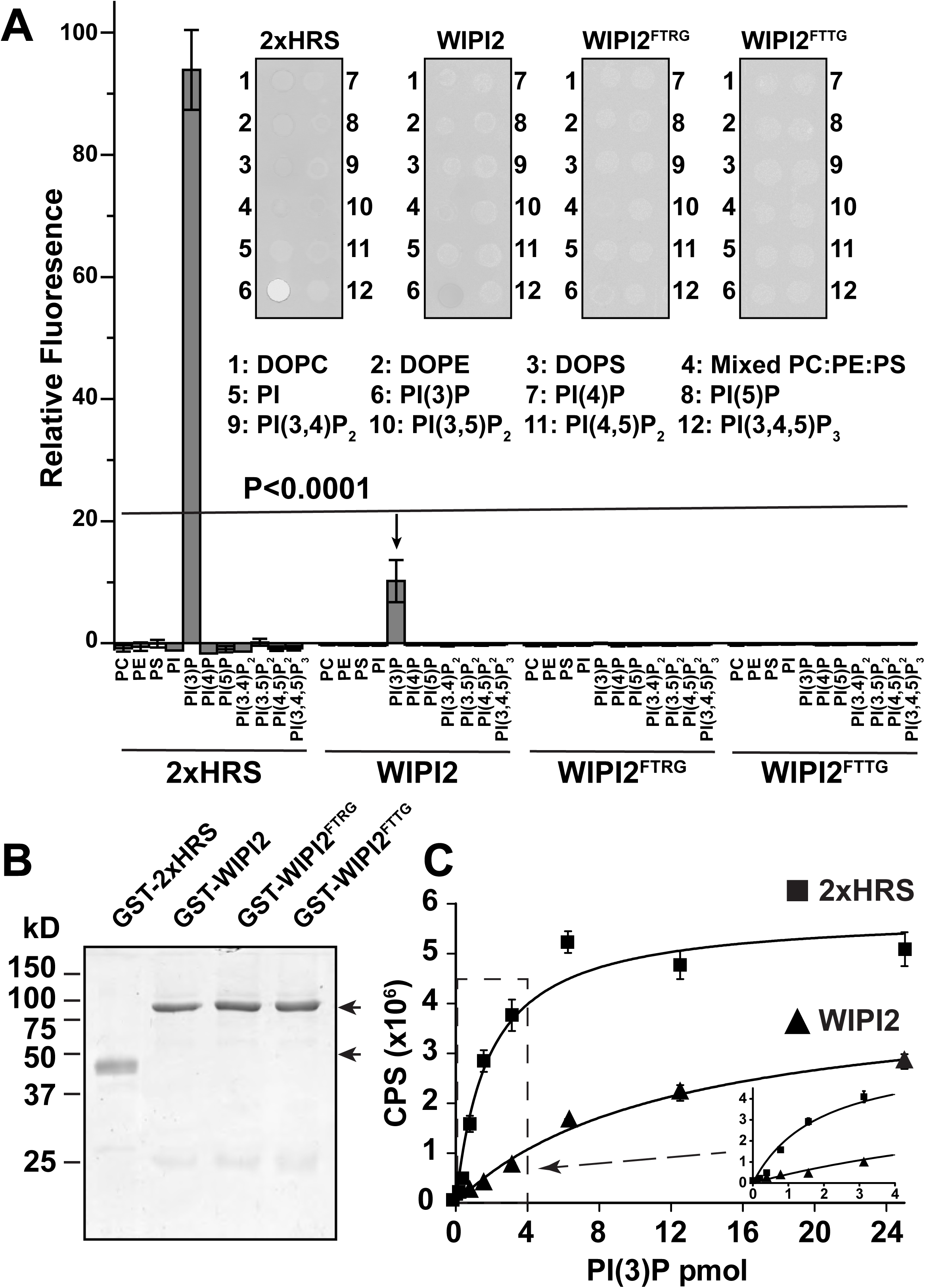
WIPI2 binds specifically to PI(3)P. (**A**) Quantification of binding of 2xHrs, WIPI2, or WIPI2 PI-binding site mutants, in overlay assays. Bars show means ± SEM from 3 experiments, each with three internal replicates. Inset, example overlay assay results. Pairwise comparisons were made by Student’s t test between signal for PI(3)P binding by WT WIPI2, and all the other samples, yielding p<0.001 for each. (**B**) Coomassie gel of purified GST-tagged probes used in (A) and (C). (**C**) Titration of PI(3)P in PI-ELISA assays. Data are fit to hyperbolic saturation binding curves. Points are means ± SEM from three experiments, each with triplicate measurements. Inset, zoomed-in view of the region shown by the dashed box.

To identify phosphoinositides in autophagosomes, hRPE1 cells were transfected with various phosphoinositide-specific probes and treated with chloroquine (CQ) to induce accumulation of autophagosomes, marked by LC3, through inhibition of lysosome fusion (Fig 8). LC3-positive structures were surrounded by large structures enriched in PI(3)P/2xHrs (Fig 8B). This arrangement has previously been described as “adjacent co-localization” (Itakura and Mizushima, 2010), and similar PI(3)P-containing ring-like structures surrounding LC3 were termed omegasomes (Axe et al, 2008). Given the resolution of optical microscopy and the fact that the signals show little true overlap, the compartments they are in must be separated by at least a few hundred nanometers. Thus, if LC3 lipidation occurs on PI(3)P-enriched membranes, it must be rapidly transported to a distinct internal compartment of the autophagosomes upon lipidation. DsRed-WIPI2 and GFP-WIPI1, in contrast to GFP-2xHrs, showed true co-localization at the resolution of confocal microscopy with many, but not all, of the LC3-positive puncta; in some cases they appeared enriched on the outside edges of the stained structures as seen for 2xHrs (Fig 8C,G). The pleckstrin homology domain from PLCδ, a probe for PI(4,5)P_2_, also partially colocalized with LC3 (Fig 8F). The PI(4,5)P_2_ localization seems to be a mixture of true co-localization, and location in adjacent membranes as seen for 2xHrs. The distinct morphologies of the compartments labeled by different PI-probes, and of the compartments containing WIPI1, WIPI2, and lipidated LC3, indicates that there are multiple distinct membrane compartments in the vicinity of the autophagosome markers. Although a previous report (Vicinanza et al, 2015) identified PI(5)P in autophagosomes, we did not detect colocalization of LC3 and DsRed-3xING2, a probe for PI(5)P, in hRPE1 cells (Fig 8E). Probes for PI(4)P and PI(3,4)P_2_ also did not co-localize with LC3 (Fig 8D,H).

**Figure 8.**
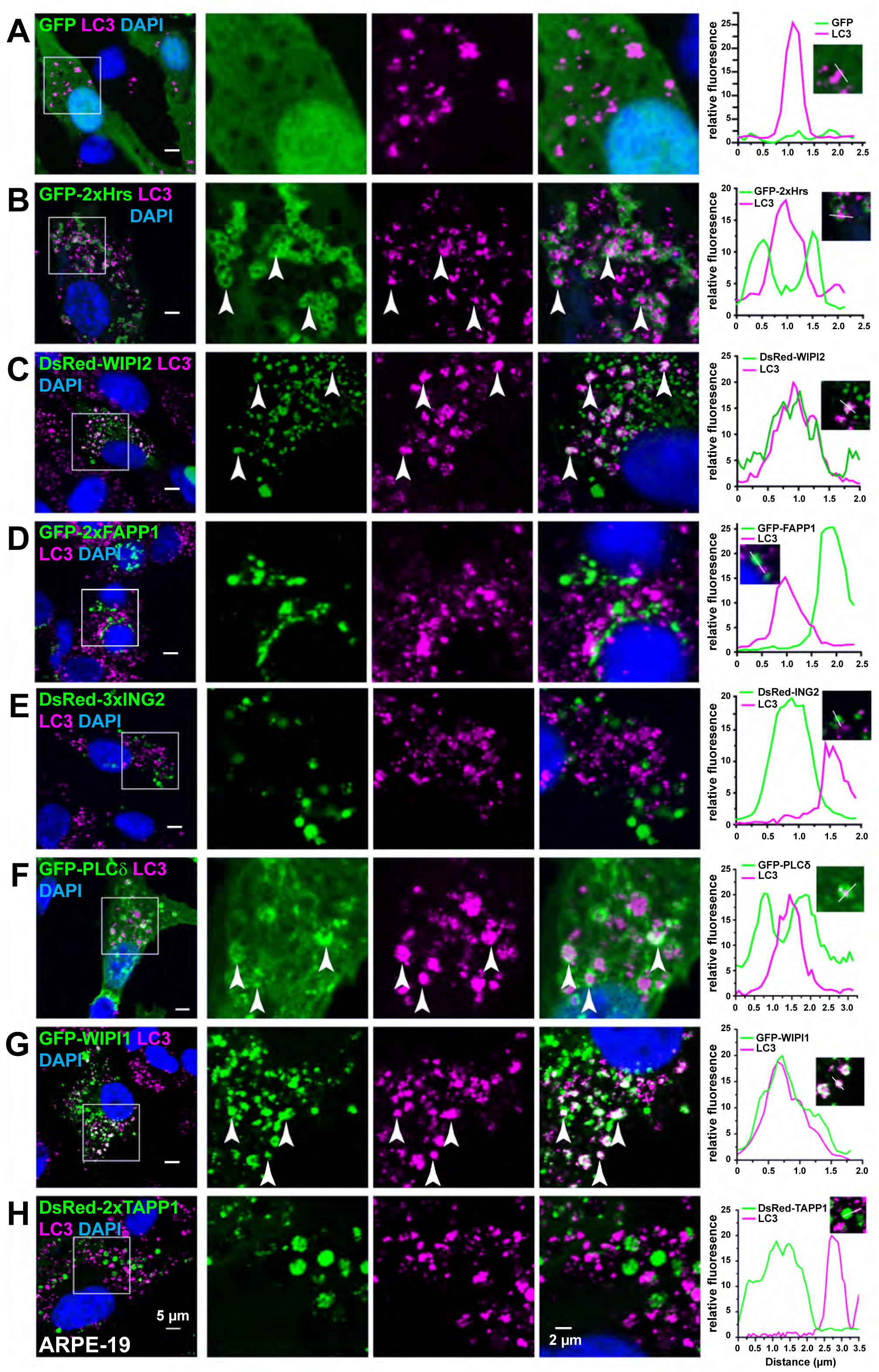
Multiple phosphoinositides and distinct membrane compartments in the vicinity of LC3-positive structures. (**A**) hRPE1 cells were transfected with phosphoinositide-binding probes, treated with CQ, and immunostained with LC3 antibody. Specific probes are: GFP-2xHrs and DsRed-WIPI2, PI(3)P; GFP-2xFAPP1, PI(4)P; DsRed-3xING2, PI(5)P; GFP-PLCδ, PI(4,5)P2; TAPP1, PI(3,4)P2.

Besides phosphoinositides, Rab11a has also been implicated in WIPI2 recruitment to autophagosome membranes (Puri et al, 2018). However, we did not detect Rab11a colocalization with WIPI2 puncta (Fig 9A). Furthermore, knockdown of Rab11a in ARPE-19 cells had no noticeable effect on WIPI2 puncta observed following CQ treatment, even after acute Vps34 inhibitor treatment (Fig 9B,C). These data indicate that whereas WIPI2 requires PI(3)P for initial recruitment to membranes, once it is bound it can maintain its membrane association in the absence of both PI(3)P and Rab11a.

**Figure 9.**
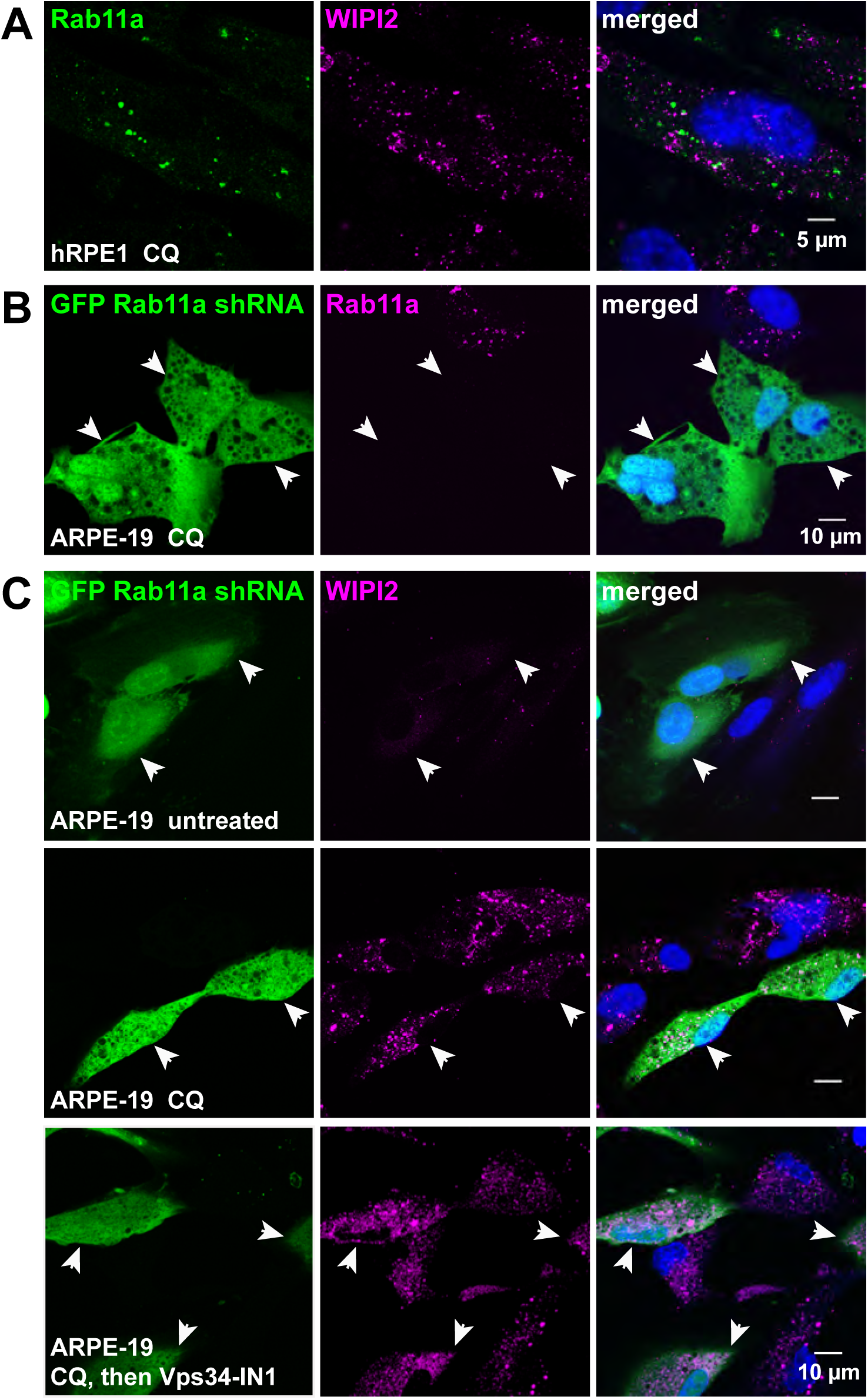
WIPI2 puncta are maintained in the absence of both PI(3)P and Rab11a. (**A**) In hRPE1 cells treated with CQ, Rab11a did not colocalize with WIPI2. (**B**) Rab11a immunostaining of ARPE-19 cells transfected with Rab11a-shRNA and treated with CQ confirmed effective knockdown. Transfected cells were identified by GFP expression from a separate expression cassette in the shRNA plasmid. (**C**) WIPI2 puncta in ARPE-19 cells were unaffected by Rab11a knockdown, in either absence or presence of acute VPS34-IN1 treatment.

Recently, Lystad et al. reported near-normal levels of LC3 lipidation in WIPI2-KO HEK cells treated with starvation or monensin, and found that under certain conditions, membrane-binding activity of ATG16L1, along with ATG3, ATG7, and ATG5-12, was sufficient to promote LC3 lipidation *in vitro* in the absence of both WIPI2 and PI(3)P (Lystad et al, 2019). In agreement with these results, ATG16L1, which formed puncta colocalizing with WIPI2 in CQ-treated control cells (Fig 10E), maintained a punctate appearance following WIPI2 knockdown with an shRNA targeting all four WIPI2 isoforms (WIPI2a-e) (Fig 10F). Furthermore, WIPI2 knockdown had no effect on CQ-induced LC3 puncta in ARPE-19, hRPE1, or HEK cells (Fig 10A-B,S5). LC3 puncta induced by starvation, bafilomycin A, or monensin were similarly unaffected by WIPI2 knockdown (Fig S6). p62-positive puncta in CQ-treated WIPI2-knockdown cells colocalized with both LC3 and Atg9 (Fig 10C-D), confirming their identity as autophagosomes. However, unlike the Vps34-specific inhibitor (Fig 9A), overnight treatment with the nonspecific kinase inhibitor wortmannin did abolish LC3 puncta in both WIPI2 knockdown cells and neighboring untransfected cells (Fig 12D), suggesting some phosphoinositide or other kinase product, but not PI(3)P or WIPI2, is required for LC3 recruitment and lipidation.

**Figure 10.**
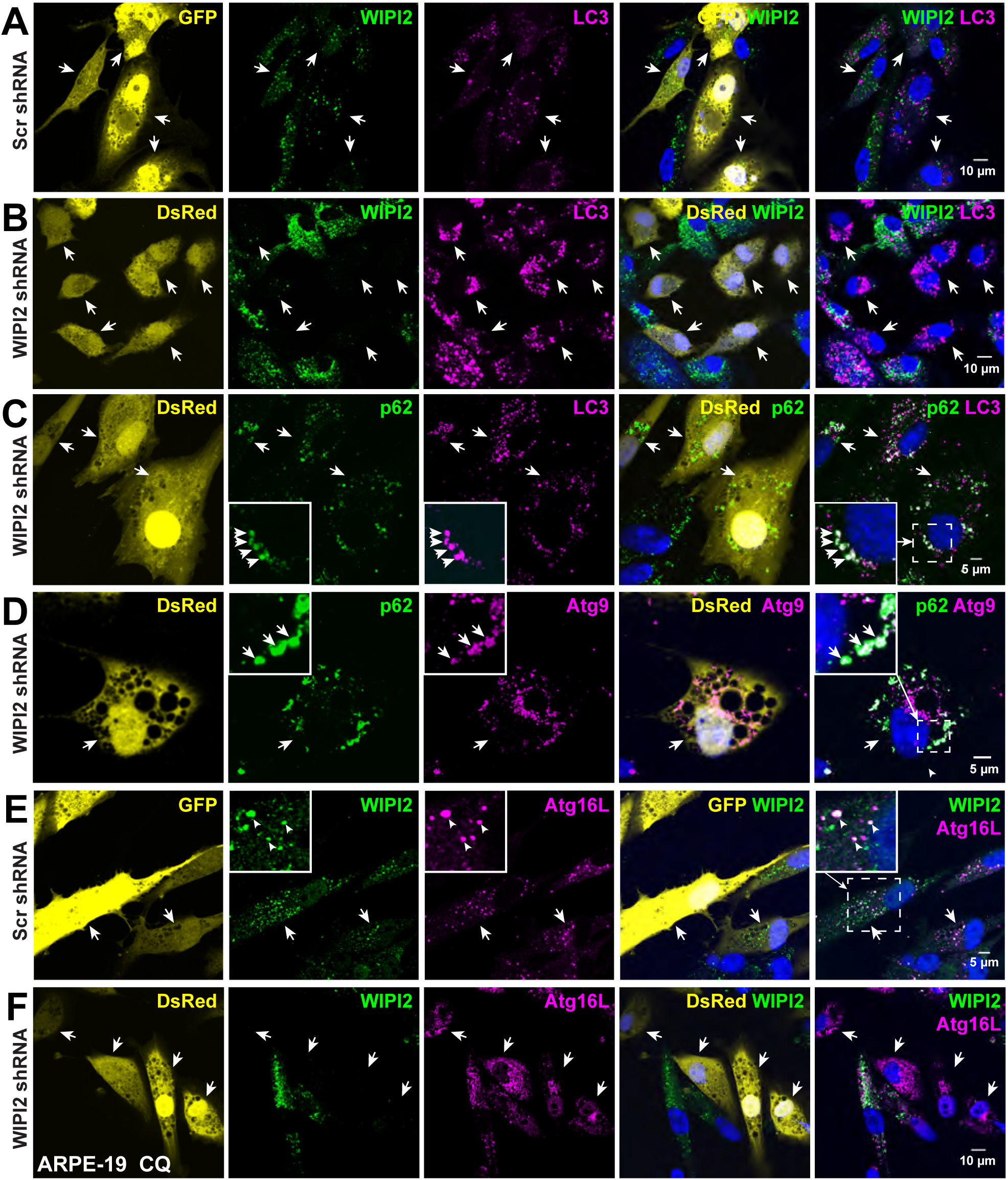
WIPI2 is dispensable for LC3 and ATG16L1 puncta formation. (**A,B**) ARPE-19 cells were transfected with scrambled (A) or WIPI2 (B) shRNA, marked by GFP or DsRed expression, respectively. WIPI2 immunostaining confirmed knockdown efficiency; LC3 puncta were unaffected. (**C,D**) Puncta formed in WIPI2 knockdown cells contain LC3, p62, and Atg9. (**E**) In ARPE-19 cells transfected with scramble shRNA control, WIPI2 puncta colocalize with Atg16L. (**F**) Atg16L1 puncta were maintained in WIPI2 knockdown cells.

## Discussion

Our results reveal an essential role for Vps34 and PI(3)P in three different membrane-recycling pathways and in maintenance of cell health in the RPE. Gross morphology defects and cell degeneration were observed in the Vps34^ΔRPE^ animals (Fig 1, 2), along with accumulation of intracellular structures identified as endocytosis, phagocytosis, and autophagy intermediates (Fig 3-5). While there are distinct roles for Vps34 and PI(3)P in each pathway, they have some features in common. In all three, PI(3)P is not needed for initiation or formation of intracellular membrane-enclosed organelles targeted for lysosomal degradation, but is absolutely essential for reaching the final step of lysosomal fusion.

Autophagy was arrested prior to lysosome fusion, evidenced by accumulation of immature autophagic structures containing p62, ATG9, and LC3, but not lysosomal marker LAMP1 (Fig 5). These observations are consistent with the autophagy defects reported in various Vps34 knockout cells and tissues (Zhou et al, 2010; Jaber et al, 2012; Bechtel et al, 2013; Reifler et al, 2014; He et al, 2016, 2019) There is no evidence that PI(3)P is needed for autophagosome formation or LC3 recruitment and lipidation in RPE, but we cannot rule out the possibility that it might be essential for them under some conditions.

WIPI2, which was identified as a PI(3)P effector protein recruited to autophagosomes in a PI(3)P dependent manner (Dooley et al, 2014; Polson et al, 2010), was also dispensable for LC3 lipidation: treatment of RPE cells with a selective Vps34 inhibitor abolished CQ-induced WIPI2 puncta, but not LC3 puncta (Fig 6); and transfection of RPE cells with WIPI2 shRNA greatly reduced WIPI2 protein levels, but did not affect LC3 puncta (Fig 10). The LC3-positive structures that accumulated in both Vps34 KO cells and CQ-treated WIPI2 knockdown cells were also positive for p62 and ATG9 (Fig 5, 10), strongly suggesting that they are autophagosomes, and many double- and multi-membrane structures were observed in Vps34 KO RPE cells by electron microscopy (Fig 5,S2). These autophagosomes formed in the absence of Vps34 could arise from nutrient deprivation due to loss of POS digestion or simply accumulate due to blocked basal autophagic flux; it is possible that different triggers for autophagy initiation may use different mechanisms for LC3 recruitment and lipidation.

Evidence for the roles of PI(3)P in autophagy is clouded by the use of a poorly-selective kinase inhibitor, wortmannin, in many studies. Consistent with previously-reported results (Itakura and Mizushima, 2010; Polson et al, 2010; Dooley et al, 2014; Fletcher et al, 2018), we observed loss of LC3 puncta following wortmannin treatment (Fig 6). However, the selective Vps34 inhibitor Vps34-IN1 was not able to abolish LC3 puncta (Fix 6), although we verified that it led to loss of PI(3)P-positive puncta (Fig. S4). These data suggest that in some cases the effect of wortmannin may be attributed to a Type I or Type II PI-3 kinase, to a Type III PI-4 kinase (Nakanishi et al., 1995), or to other off-target effects, *e.g.* on a protein kinase as previously documented (Liu et al, 2007). The ATG12-ATG5-ATG16L1 complex is required for LC3 lipidation (Mizushima et al, 2001, 2003; Fujita et al, 2008). Though WIPI2 is a direct binding partner of ATG16L1 (Dooley et al, 2014), and in some cases WIPI2 potentiates LC3 lipidation (Polson et al, 2010; Dooley et al, 2014; Lystad et al, 2019; Stavoe et al, 2019), it is not required for LC3 lipidation induced by either starvation or monensin in HEK-293 cells (Lystad et al, 2019). ATG16L1 was previously shown to possess its own membrane-binding activity (Fletcher et al, 2018; Lystad et al, 2019). We also observed ATG16L1 puncta in CQ-treated WIPI2 knockdown RPE cells (Fig 10), supporting a mechanism utilizing direct membrane binding of ATG16L1. However, the mechanism of its recruitment specifically to the appropriate membranes in the absence of WIPI2 is unknown.

The association of LC3 with the arrested autophagosomes indicates that PI(3)P is dispensable for LC3 lipidation in autophagosome formation but is required for a downstream step leading to lysosome fusion. In some cells treated with Bafilomycin A, which blocks lysosome acidification and autophagosome-lysosome fusion, LC3 and lysosome markers colocalize (Fig S6; Mauthe et al, 2018; but see Mauvezin et al., 2015), suggesting interaction of the two compartments. The lack of co-localization of lysosomes and autophagosomes in Vps34 KO cells (Fig 5) suggests, in contrast, a defect in transport of autophagosomes and/or lysosomes. Microtubule-based motors are required for autophagosome transport (Kochl et al 2006; Kimura et al, 2008), and a similar lack of lysosome co-localization was observed when cells were treated with microtubule-disrupting drugs (Kochl et al, 2006). PI(3)P-binding proteins have also been implicated in autophagosome transport and autophagolysosome fusion (Pankiv et al, 2010; Chen et al, 2012; Olsvik et al, 2015), consistent with defects in autophagosome maturation observed in the Vps34 KO cells. A critical role has been proposed for the PI(3)P-binding PIP-5 kinase, PIKfyve, and its product PI(3,5)P_2_ in endosome maturation (de Lartique et al. 2009), and PI(3)P has been reported to be important for lysosome positioning (Hong et al., 2017).

It was recently reported that non-lipidated LC3 can form aggregates in Atg9 and Atg16L1 KO HeLa cells, which also colocalized with p62 and ubiquitin, raising the possibility of erroneous interpretation of LC3 puncta (Runwal et al, 2019). In the case of Vps34 KO, however, robust LC3 lipidation has been confirmed by Western blot, by us and others (He et al, 2016; Jaber et al, 2012; Bechtel et al, 2013), and LC3 puncta colocalize with Atg9 (Fig 5). The LC3 aggregates in Atg9 and Atg16L1 KO cells were significantly different in size and number from those in WT cells (Runwal et al, 2019), whereas the size and number of LC3 puncta in WIPI2 knockdown cells was not noticeably different from untransfected cells in the same field (Fig 10), and likewise the LC3 puncta in Vps34-IN1 treated cells were similar in size and number to those in control cells (Fig 6).

LC3 is also recruited to phagosome membranes in RPE (Fig 4) (Frost et al, 2015). In Vps34 KO RPE, POS-containing phagosomes accumulated prior to association with lysosomes and fusion; however, unlike autophagosomes, the accumulated phagosomes were LC3-negative, indicating that PI(3)P is not needed for engulfment but is critical for LC3 recruitment and, by inference, lipidation (Fig 4). Previous reports have indicated divergent mechanisms of ATG16L1 recruitment to phagosomes and autophagosomes in other cell types. LC3 recruitment to phagosomes was found to be PI(3)P independent (Florey et al, 2015; Fletcher et al, 2018), and the membrane-binding C-terminal domain of ATG16L1 was required for LC3 lipidation on phagosomes, but not autophagosomes (Fletcher et al, 2018; Rai et al, 2019), suggesting that direct membrane binding of ATG16L1 is employed on phagosome membranes, consistent with PI(3)P being dispensable for ATG16L1 recruitment to phagosomes. Our finding that PI(3)P is required for LC3 lipidation on POS-containing phagosomes in the RPE is in direct contrast to these reports, and suggests that RPE cells utilize a distinct adaptation of the LAP pathway.

The maturation pathway of POS phagosomes in the RPE is poorly understood. Possibilities might include similar pathways downstream of LC3 lipidation as found in autophagy, or interactions with endosomes (Wavre-Shapton et al, 2014); however we did not detect autophagy markers or endosome markers colocalized with POS phagosomes, nor did we detect preferential degradation of the RHO C-terminus which has been associated with endosome fusion (Wavre-Shapton et al, 2014).

PI(3)P clearly plays a critical role in endosome processing as well. In resting RPE cells, PI(3)P strongly co-localizes with endosome markers Rab5 and Rab7 (Fig S1), and Rab7-positive late endosomes accumulated in the Vps34 KO cells (Fig 5). The KO cells also have drastically altered levels and distribution of the early endosome marker EEA1 (Fig 1). These observations point to a critical need for PI(3)P in endosome processing and particularly in endosome-lysosome fusion.

Ultimately, our results show that phagocytosis, autophagy, and endocytosis in the RPE are severely impaired in the absence of PI(3)P, although the affected steps differ. The results presented here highlight the critical importance of PI(3)P for multiple types of degradative membrane processing, and the consequence of their failure on cell health. They also highlight the fact that there are multiple and possibly redundant pathways for recruitment of LC3 to membranes, and that different stimuli and different cell types may mobilize different pathways.

## Materials and Methods

### Animals

All animal studies were conducted according to NIH guidelines and approved by the Institutional Animal Care and Use Committee at Baylor College of Medicine. Wild-type C57BL/6J, B6 albino (B6(Cg)-Tyrc-2J/J), and transgenic mice that express Cre recombinase under control of the human bestrophin-1 (*BEST1*) promoter (C57BL/6-Tg(*BEST1-cre*)1Jdun/J) (Iacovelli et al, 2011) were purchased from Jackson lab (Bar Harbor, ME). CD-1 IGS mice were purchased from Charles River (Wilmington, MA). Vps34 floxed mice which have lox P sites flanking exons 17 and 18 (the ATP binding domain) (Zhou et al, 2010) were a gift from Fan Wang (Duke University). Vps34 floxed (Vps34^fl/fl^) mice were back-crossed to C57BL/6 for at least 6 generations and bred with the *BEST1*-Cre mice to generate RPE-specific conditional functional Vps34 knockout mice (Vps34^ΔRPE^). The B6 mice were crossed with Vps34 knockout mice to obtain albino Vps34^fl/fl^ and Vps34^ΔRPE^. Vps34 knockout mice that were also heterozygous for the human rhodopsin-GFP allele at the mouse rhodopsin locus (Vps34^ΔRPE^;Rho^+/hrhoG(H)^) were generated by crossing Vps34^ΔRPE^ mice with human rhodopsin-GFP knock-in mice (hrhoG(H)) containing a lox H site in the 5’ non-coding region to reduce the human rhodopsin-GFP expression (Chan et al, 2004).

Mice used for tissue samples were euthanized by CO_2_ inhalation prior to dissection following American Association for Laboratory Animal Science protocols. All the mice were maintained in a 12/12-hour light (400 Lux)/dark cycle.

Tail DNA was used for Vps34 allele genotyping as described (Zhou et al, 2010). PCR primers 5′-ATGCCCAAGAAGAAGAGGAAGGTGTCC -3′ and 5′-TGGCCCAAATGTTGCTGGATAGTTTTTA -3′ were used for PCR analysis of the *BEST1*-Cre transgene (Iacovelli et al, 2011). The absence of Pde6b^rd1^ was confirmed by PCR (Gimenez & Montoliu, 2001); absence of Crb1^rd8^ was confirmed by PCR and sequencing of the relevent Crb1 region (Mattapallil et al, 2012).

### Cell culture and treatment

ARPE-19 (CRL-2302; ATCC, Manassas, VA) and hTERT RPE-1 (CRL-4000; ATCC) were cultured in Corning DMEM/Ham’s F-12 1:1 mixture (Fisher Scientific, Waltham, MA) with 10% FBS (Sigma, St. Louis, MO) and were maintained at 37°C in 5% CO_2_. ARPE-19 cells used in phagocytosis assays were cultured in as above but with 10% heat-inactivated FBS. The ARPE-19 and hTERT RPE-1 RPE cells used in all experiments were cultured for less than 20 passages. HEK-293 cells were from ATCC (#CRL-1573) and cultured in DMEM with 10% FBS. Sf9 insect cells were used to express wild type and mutant mouse WIPI2. Sf9 cells were cultured in flasks with Lonza Insect Xpress medium (Fisher Scientific) containing 10% heat inactivated FBS, 50 units/ml penicillin, and 50 µg/ml streptomycin at 27°C.

Drug treatments were: 25 µg/ml chloroquine diphosphate salt (C6628), 100 nM bafilomycin A1 (SML1661), 100 µM monensin sodium salt (M5273), and 10 µM wortmannin (12-338) that were purchased from MilliporeSigma or 20 µM Vps34IN1 (S7980; Selleckchem, Houston, TX). Overnight drug treatments were for 12 hours, and acute treatments were for 90 min or as indicated. Control cells were treated with an equal volume of DMSO.

For the transfection experiments, RPE or HEK cells were seeded in 24-well plates with coverslips. 0.4 µg DNA, 2 µl ViaFect™ (Promega, Madison, WI) and 40 µl Gibco Opti-MEM reduced serum media (Fisher Scientific) were mixed well and incubated at room temperature for 15 min, then added into each well containing 0.5 ml culture medium. For shRNA knock down, cultured cells were transfected with 0.4 μg shRNA-expressing plasmid DNA with ViaFect and cultured for 5 days before immunostaining.

### Constructs

Hepatocyte growth factor-regulated tyrosine kinase substrate (Hrs, GenBank: AAH03239.1) phosphoinositide-binding domain (amino acids 147-223) was cloned from purified C57BL/6 mouse kidney mRNA as described (He et al, 2016). Mouse tandem PH domain-containing protein-1 (TAPP1) PH domain cDNA (amino acids 181-300) (Dowler et al, 2000) was a gift from Dario R. Alessi (University of Dundee, Scotland, U.K.); phospholipase C δ1 (PLCδ) PH domain (amino acids 1-175) (Stauffer et al, 1998) was a gift from Tobias Meyer (Stanford University, CA); The sequence of four-phosphate-adaptor protein 1 (FAPP1) PH domain was identical to amino acids 1-100 of human FAPP1 (GenBank: AF286162.1); inhibitor of growth protein-2 (ING2) cDNA was a gift from Curtis Harris (Addgene plasmid # 13294) (Pedeux et al, 2005); and WD repeat domain phosphoinositide-interacting protein 1 (WIPI1) cDNA was a gift from Noboru Mizushima (Addgene plasmid # 38272) (Itakura & Mizushima, 2010). Mouse WD repeat domain phosphoinositide-interacting protein 2 (WIPI2) cDNA was cloned from mouse retina phage cDNA (ʎZAPII) library (a gift from Wolfgang Baehr, University of Utah) using primers 5’-ATGAACCTGGCGAGCCAGAGCGGAGAGGCC-3’ and 5’-ACAGCGAACATCCTCCCATGATTCTCCGGACTGAC-3’, and the sequence was confirmed as identical to NM_178398.4. Two tandem copies of Hrs, FAPP1, or TAPP1, three tandem copies of ING2, or one copy of PLCδ phosphoinositide-binding domains, or WIPI1 or WIPI2 cDNA were cloned with N-terminal EGFP or DsRed fusions in pCAG vector (a gift from Connie Cepko (Addgene plasmid #11150)) (Matsuda & Cepko, 2004) using EcoRI and NotI sites. All constructs were verified by sequencing.

For purification in E.coli, a C-terminal 1D4 epitope tag (TETSQVAPA) was added to 2xHrs by PCR, then sub-cloned into the pGEX vector (GE Life Sciences), thereby also adding an N-terminal GST tag. Residues in the 4-amino acid phosphoinositide binding sequence FRRG in mouse WIPI2 were mutated to FTRG or FTTG (Baskaran et al, 2012; Liang et al, 2019) by PCR site-directed mutagenesis. WT mouse WIPI2, WIPI2^FTRG^, and WIPI2^FTTG^ with N-terminal GST fusions and C-terminal 1D4 tags were cloned into BamHI and NotI sites in pFb1 vector (Thermo).

For shRNA constructs, vectors pBS-U6-EGFP and pBS-U6-DsRed were generated by subcloning the murine U6 promoter from pCS6/U6 (a gift from Connie Cepko (Addgene plasmid # 15173) (Harpavat & Cepko, 2006)) to pBluescript II KS(-) using BssHI sites, then inserting a downstream EGFP or DsRed expression cassette containing a CMV promoter and BGH polyA signal, derived from pCDNA3.1. Target sequences were chosen for screening based on scores from the Broad Institute GPP Web Portal, with preference for targets starting with G, allowing authentic N-terminal transcription beginning at the residual G of the ApaI site (Shukla et al, 2007). Annealed oligos were ligated into pBS-U6 vectors digested with ApaI (blunted) and EcoRI, as described (Matsuda & Cepko, 2004). shRNA constructs were screened for effectiveness by transfection and immunostaining for the target protein, and comparing transfected cells, marked by GFP or DsRed fluorescence, with untransfected cells in the same field. Oligos for shRNAs in this study were (target sequence underlined): Rab11a, 5’-

TATCATGCTTGTGGGCAATAACTCGAGTTATTGCCCACAAGCATGATACTTTTTG-3’ and 5’-

AATTCAAAAAGTATCATGCTTGTGGGCAATAACTCGAGTTATTGCCCACAAG

<u>CATGATA-3’</u>; WIPI2, 5’-

CTGTCAATCAACAACGACAACTCGAGTTGTCGTTGTTGATTGACAGCTTTTTG-

3’ and 5’-

AATTCAAAAAGCTGTCAATCAACAACGACAACTCGAGTTGTCGTTGTTGATT

<u>GACAG</u>-3’ or 5’-

AGACCGTGCACATCTTCAAACTCGAGTTTGAAGATGTGCACGGTCTCTTTTTG

-3’ and 5’-

AATTCAAAAAGAGACCGTGCACATCTTCAAACTCGAGTTTGAAGATGTGCAC

<u>GGTCT</u>-3’. Scramble control sequence was from (Sarbassov et al, 2005): 5’-

CCTAAGGTTAAGTCGCCCTCGCTCGAGCGAGGGCGACTTAACCTTAGGCTTTTTG-3’ and 5’-

AATTCAAAAAGCCTAAGGTTAAGTCGCCCTCGCTCGAGCGAGGGCGACTTAA

CCTTAGG-3’.

### Expression and purification of phosphoinositide binding proteins

GST-2xHrs-1D4 was expressed in BL21(DE3)pLysS E. coli (MilliporeSigma, Burlington, MA) at room temperature for 4 hours following induction with 1 mM isopropyl β-D-1-thiogalactopyranoside. The cells were harvested and lysed in 40 mM Tris, pH 7.6, 0.2 M NaCl, 5 mM DTT, 1% triton X-100, phenylmethanesulfonyl fluoride (PMSF) and cOmplete™ Protease Inhibitor Cocktail (Roche). Protein purification was performed with Glutathione Sepharose 4 Fast Flow (GE Healthcare) following the manufacturer’s protocol. Before elution, the detergent was washed out with 40 mM Tris, pH 7.6, 0.2 M NaCl, 5 mM DTT, then protein was eluted with 40 mM Tris, pH 7.5 with 40 mM reduced L-Glutathione (Sigma). Purified proteins were dialyzed against 20 mM Tris pH 7.5, 100 mM NaCl, and PMSF, then concentrated, supplemented with 50% glycerol and PMSF, and stored at -20°C.

GST-WIPI2-1D4, GST-WIPI2^FTRG^-1D4 and GST-WIPI2^FTTG^-1D4 were expressed in Sf9 insect cells. Recombinant baculovirus was generated using the Bac-to-Bac baculovirus expression system (Fisher Scientific) and was produced in Sf9 cells. Passage 3 or 4 virus was used to infect Sf9 cells for protein production. Cells in a 150 cm^2^ flask were adsorbed with 0.5 ml of virus and 2-3 ml Insect Xpress medium for 1 hour at room temperature with frequent mixing, followed by addition of medium. Cells from 5-6 flasks were harvested at 50-60 hrs post-infection. The cell pellets were lysed by sonication in 40 mM Tris, pH 7.6, 0.2 M NaCl, 5 mM DTT, PMSF, and cOmplete™ Protease Inhibitor Cocktail. Protein purification was performed with Glutathione Sepharose as described above.

### Phosphoinositide binding assays

For phosphoinositide overlay assays (Furutani et al, 2006), synthetic dipalmitoyl diC16 PI, PI(3)P, PI(4)P, PI(5)P, PI(3,4)P_2_, PI(4,5)P_2_, PI(3,5)P_2_, and PIP3 standards were purchased from Echelon Biosciences (Salt Lake City, UT). 2 µl of 50 pmol/µl phosphoinositide containing 3.75 nmol mixed phospholipids (PC, PE, and PS in 45:35:15 molar ratio) (Echelon Biosciences and Avanti Polar Lipids, Alabaster, Alabama) in methanol:chloroform (9:1 v/v) solvent was loaded on to 0.45 µm supported nitrocellulose membrane (GVS, Sanford, ME). The membrane was air-dried for 1 hour at room temperature, then blocked with 5% non-fat milk and 3% BSA in PBS for 3 hours at room temperature, incubated with 40 nM GST-2xHrs-1D4 or GST-WIPI2-1D4 protein in PBS with 3% BSA at 4°C overnight. The membrane was washed with TBST and incubated with 1D4 antibody (1 μg/ml in PBS with 3% BSA) at room temperature for 3 hours, followed by goat anti-mouse IRDye 800CW (LI-COR, Lincoln, NE) diluted 1:10,000 in PBS with 3% BSA for 40 mins. Membranes was imaged and analyzed with an Odyssey infrared imaging system (LI-COR). The samples were triplicate in each assay and the assays were repeated at least 3 times.

The phosphoinositide-ELISA assay was performed as described (He et al, 2016), with some modifications. Standards containing 0 to 25 pmol PI(3)P and 5 nmol total PC, PE, PS in 45:35:15 molar ratio in 40 µl methanol:chloroform (9:1 v/v) were added into each well of a PolySorp 96-well plate (Thermo Fisher). All samples were performed in triplicate. Lipids were air-dried, then dried under vacuum overnight. The plate was blocked with 5% BSA (Sigma) in PBS for 3 hours at room temperature, then incubated with 100 µl/well of 40 nM purified GST-2xHrs-1D4 or GST-WIPI2-1D4 in PBS with 3% BSA at 4°C overnight. The plate was incubated with 1D4 antibody (1 μg/ml in PBS with 3% BSA) at room temperature for 3 hours, washed with PBS 6 times, then incubated with goat anti-mouse-HRP (Thermo) diluted 1:3000 in PBS with 3% BSA at room temperature for 1 hour. After washing with PBS, 100 µl of SuperSignal ELISA Femto Maximum Sensitivity Substrate (Thermo) were added to each well, and after ∼1 min, photons were detected using a Victor3 Multilabel Plate Counter (Perkin Elmer). The limit of detection was about 0.1-0.2 pmol/well. The assays were repeated more than 3 times. Data were analyzed and fit to hyperbolic saturation binding curves using GraphPad Prism and Origin software.

### Sub-retinal injection and electroporation

pCAG-GFP or pCAG-GFP-2xHrs plasmid DNA (2 mg/ml) was introduced to the RPE by sub-retinal injection and electroporation of P0-P1 WT CD1, albino Vps34^fl/fl^, or albino Vps34^ΔRPE^ mice as described (Agosto et al, 2018; Matsuda & Cepko 2004, 2008), except that the electrodes were reversed such that the RPE, rather than the neural retina, was electroporated. Mouse pups were anesthetized, and the future edge of the eyelid was opened with an incision. A pilot hole was made with a 30-G beveled needle, a 33-G blunt needle was positioned in the subretinal space, and ∼450 nl of plasmid DNA in PBS with 0.1% Fast Green dye was injected using a UMP3 Microsyringe Injector and Micro4 Controller (World Precision Instruments). Five square wave pulses (80 V, 50 ms) were applied with an ECM 830 square wave electroporator (BTX Harvard Apparatus) and custom tweezer-type electrodes, with the negative electrode on the injected eye. Injected eyes were fixed 8 weeks later.

### Immunofluorescence staining

The following antibodies and reagents were used for immunostaining: rabbit anti-Cre (#69050), and mouse anti-ubiquitin (#04-263) were from EMD Millipore; rabbit anti-GRP78/BiP (#PA1-014A), rabbit anti-TGN46 (#PA5-23068), Armenian hamster anti-Atg9 (#MA1-149), rabbit anti-ZO-1/TJP1 (#402200) and Alexa Fluor 647 phalloidin (#A22287) were from Thermo Fisher; mouse anti-GM130 was from BD Biosciences (#610822); rabbit anti-Rab5 (#3547, C8B1), rabbit anti-Rab7 (#9367, D95F2), rabbit anti-LC3A/B (#12741) and rabbit anti-EEA1 (#3288S, C45B10) were from Cell Signaling; mouse anti-LC3 (#AM1800a) was from Abgent; guinea pig anti-p62 (#03-GP62-C) was from American Research Products; purified monoclonal 1D4 (Molday & MacKenzie, 1983) was prepared in-house from hybridoma culture medium as described (Agosto et al, 2014); mouse anti-LAMP2 (#ab25631), rabbit anti-Rab11a (#ab128913), rabbit anti-Atg9 (#PA1-16993) and mouse anti-RPE65 (#MA1-16578) were from Abcam; mouse anti-LAMP1 (##428017) was from Calbiochem; mouse anti-Atg16L (#M150) and rabbit anti-Atg16L (#PM040) were from MBL. Donkey secondary antibodies conjugated with Alexa dyes (#A21206, #A21206, #A31572, # A21202, #A31571 and #A32773) were from Thermo.

Albino mouse eyes were excised and rinsed with BPS. The eyes were fixed with 4% paraformaldehyde in PBS for 45 min at room temperature then cryoprotected with 26% sucrose in PBS at 4°C overnight and embedded in optimum cutting temperature (O.C.T.) compound for sectioning. For flat-mount eyecup preparations, eyes were fixed with 4% paraformaldehyde in PBS for 45 min at room temperature, then washed with PBS 5-6 times; the cornea, iris, lens, and retina were removed, and eight symmetric radial cuts were made. RPE flat-mounts or eyecup sections were washed with PBS, then incubated in blocking solution (10% donkey serum, 5% BSA, 0.4% fish gelatin, and 0.4% triton X-100 in PBS) at room temperature for 1 hour. The samples were incubated with primary antibodies diluted 1:50-1:100 with blocking solution at room temperature overnight.

After washing with PBS, the samples were incubated with secondary antibody conjugated with 2 µg/ml Alexa dye (Thermo) and 300 nM DAPI (Sigma) in blocking solution for 1 hour at room temperature, washed with PBS and mounted with Vectashield anti-fade reagent (Vector Laboratories).

Cultured cells on coverslips were fixed with 4% paraformaldehyde in PBS for 15 min at room temperature, then washed with PBS and incubated with blocking solution at room temperature for 0.5 hour. Coverslips were immunolabeled for 1 hr at room temperature with primary antibodies diluted 1:50-1:100 in blocking solution, followed by secondary antibodies and DAPI as above, and mounted with Prolong Gold (Thermo).

### Fluorescence microscopy and live-cell imaging

Samples were imaged with a Leica TCS SP5 confocal microscope using a Leica 63x oil immersion objective (HCX PL APO CS 63.0x, aperture1.40 oil) with lasers: diode 405, argon 488, HeNe 543, and HeNe 633. ImageJ was used for converting the original images to tiff files and contrast and brightness were adjusted in Photoshop (Adobe).

Tiled images of flat-mount RPE were acquired with the ZEISS Apotome2 Microscope system (Zeiss, Oberkochen, Germany) with 40x water immersion objective. Each tile was 1024 x 1024 with pixel size 6.45 x 6.45 µm, and ∼350-400 tiles were stitched together using ZEN 2 (blue edition) software (Zeiss). Raw images scaled to 3.23 μm/pixel were used for automated detection of p62 puncta using the “MorphologicalComponents” function in Mathematica (Wolfram), with the “ConvexHull” method and background value of 0.3. Very large features (containing ≥ 500 px), resulting from imaging artifacts at the edges of the tissue, were omitted from the data set. Flat-mount areas were determined in ImageJ from regions of interest drawn manually along the borders of the tissue.

For live-cell imaging, hRPE1 cell were seeded in Lab-TeK II 8-chambered coverglass (Nunc, Rochester, NY) and transfected with GFP-2xHrs or GFP-WIPI2. The transfected cells were treated with 25 µg/ml chloroquine for 12 hours, then recorded at room temperature with Leica TCS SP5 confocal microscope using a Leica 63x oil immersion objective. The media was replaced with fresh DMEM/Ham’s F-12 (without phenol red) containing 25 µg/ml chloroquine, and 20 μM Vps34-IN1 or an equal volume of DMSO, and cells were imaged at 30 second intervals for 90 min.

Quantification of fluorescence intensities was performed using ImageJ software. The total fluorescence intensity, minus background fluorescence, was measured for three ROIs in each time-lapse, and normalized to the intensity in the first frame.

### In vitro phagocytosis assay

POS were purified from 2-month-old heterozygous Rho^+/hrhoG(H)^ mice as described (Wensel & Gilliam, 2015). The mice were dark-adapted overnight, and retinas were collected into 300 µl of 8% OptiPrep (Sigma) in Ringers solution, vortexed at low speed for 2 min at room temperature under dim red light, and centrifuged at 400 x g at room temperature for 2 min. The supernatant containing mouse POS was collected and the retina pellet was extracted an additional 3-4 times with 8% Optiprep. The crude POS were loaded onto the top of a stepwise OptiPrep gradient (10%, 15%, 20%, 25% and 30%), and the gradient was centrifuged for 60 min at 24,700 × g at 4 °C using a TLS-55 swinging bucket rotor (Beckman Coulter). Purified mouse POS were resuspended in DMEM/Ham’s F-12 containing 2.5% sucrose. The concentration of the POS particles was estimated by Z1 Counter® Particle Counter (Beckman Coulter), and stored in the dark at -80°C.

The phagocytosis assay was based on previously-reported protocol (Mazzoni et al, 2019), with modifications. POS were treated with 1 µM MFG-E8 and 0.1 µM Gas6 (R&D Systems) in DMEM/Ham’s F-12 medium at room temperature for 30 min. ARPE-19 cells cultured on coverslips in a 24-well plate were treated with 20 µM Vps34-IN1 or the same volume of DMSO for 90 min, then were challenged with ∼10-20 treated POS particles per cell for 2 hours at 37°C in a CO_2_ incubator. The phagocytosis assay was terminated by washing the cells with ice-cold PBS-CM (PBS containing 1 mM MgCl_2_ and 0.2 mM CaCl_2_) 4-5 times. The cells were fixed with 4% paraformaldehyde in PBS for 15 min at room temperature, blocked with 1% BSA in PBS-CM for 10 min, then labeled with 1D4 antibody diluted to 1 µg/ml in 1% BSA in PBS-CM for 25 min at room temperature, followed by 2 µg/ml donkey anti-mouse Alexa Fluor 647 in 1% BSA and PBS-CM at room temperature for 25 min. After washing, the cells were blocked again with blocking solution at room temperature for 0.5 hour, then incubated with rabbit anti-LC3A/B at 1:100 dilution in blocking solution at room temperature for 1 hour, followed by donkey anti-rabbit Alexa Fluor 555, 2 µg/ml, and 300 nM DAPI in blocking solution for 1 hour at room temperature. The coverslips were washed 5 times with PBS-CM and mounted with Prolong Gold.

### Transmission electron microscopy

was performed as described (He et al, 2016). Eye cups were fixed with 3% paraformaldehyde and 3% glutaraldehyde solution in 0.1M cacodylate buffer, pH 7.4 (CB) for 1-3 days at 4°C. Eyecups were washed, post-fixed with 1% OsO4 in 0.1M CB for 1 hour, and dehydrated by sequential incubation in 30%, 50%, 70%, 85%, 90%, and 100% ethanol. The tissue was incubated with several changes of acetone and gradually infiltrated with 2:1, 1:1, and 1:2 acetone to resin (Embed-812, Electron Microscopy Sciences, Hatfield, PA) mixtures, followed by pure resin for 48 h. Resin blocks were polymerized at 60°C for 24-48 hours. 80-90nm sections were cut on a Leica Ultracut UCT microtome, collected on 50 or 75 mesh grids and stained with 2% uranyl acetate and Reynold’s lead citrate. Images were taken on a Zeiss transmission electron microscope equipped with a CCD camera (Advanced Microscopy Techniques Corp., Woburn, MA).

## Supplementary Material

The Supplementary Material consists of six supplemental figures, Figs. S1-S6.

## Acknowledgments

We thank Dr. Yan Chen (The Dean McGee Eye Institute, University of Oklahoma) for helpful suggestions and technical support. We thank Kailasam Lavanya for maintaining the mouse lines and protein expression. Supported by National Institutes of Health grants R01-EY025218 and R01-EY026545 from the National Eye Institute, and by the Welch Foundation, Q0035 (TGW). The EM facilities are supported by Core Grant for Vision Research P30-EY002520 from the National Eye Institute. The authors declare no competing financial interests.

## Author contributions

Feng He, Theodore G Wensel and Melina A. Agosto designed experiments, analyzed data and wrote the manuscript. Feng He, Ralph M Nichols, Melina A Agosto, performed experiments.

---

**Figure S1 (related to Figure 1).**
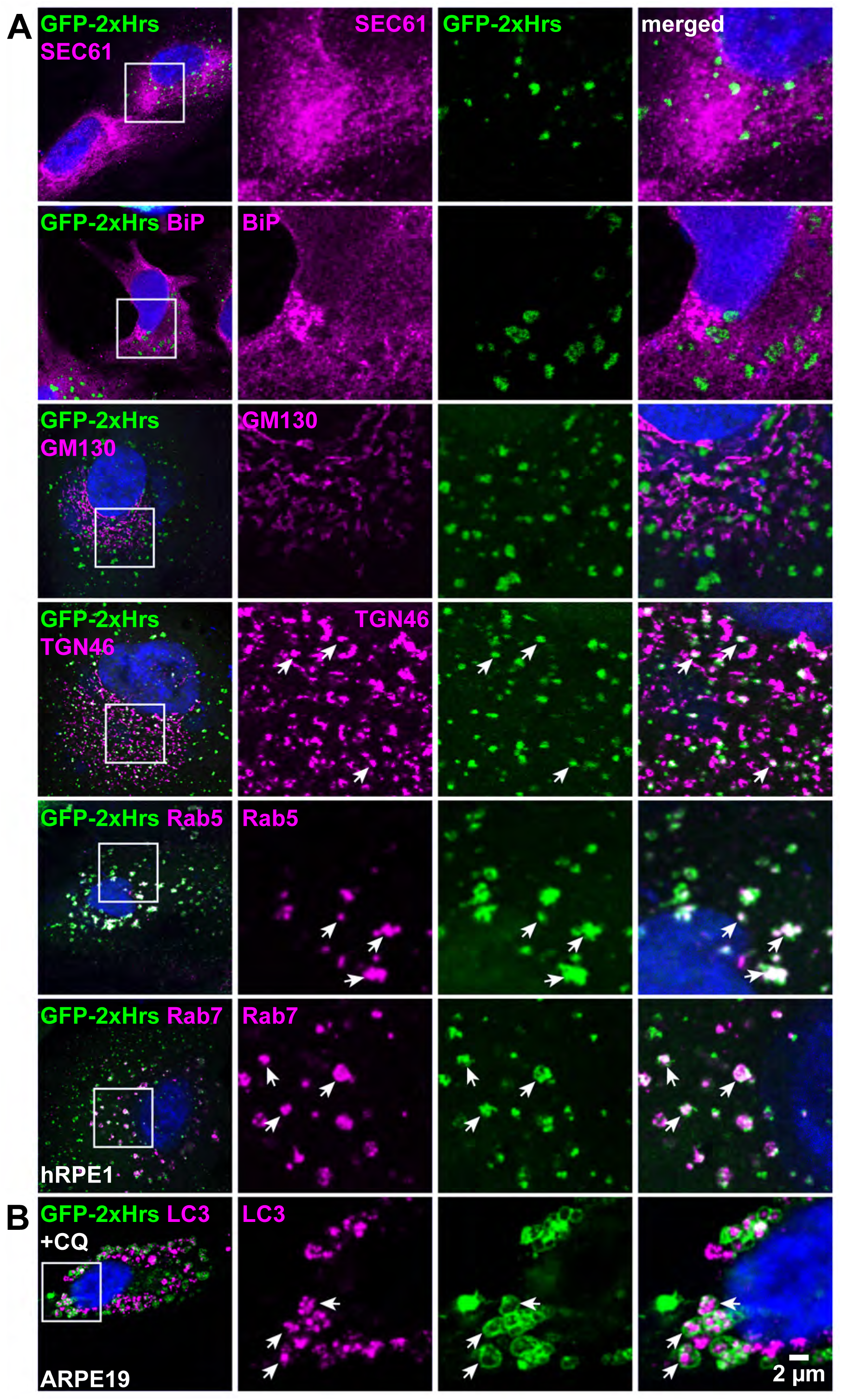
Localization of PI(3)P in ARPE-19 cells. (**A**) hRPE1 cells were transfected with GFP-2xHrs and ER marker (DsRed-Sec61β) or immunostained for ER marker BiP, *cis*-Golgi marker GM130, *trans*-Golgi marker TGN46, or endosomal markers Rab5 and Rab7. (**B**) ARPE-19 cells were transfected with GFP-2xHrs, treated with CQ, and immunostained for LC3.

**Figure S2 (Related to Figure 2).**
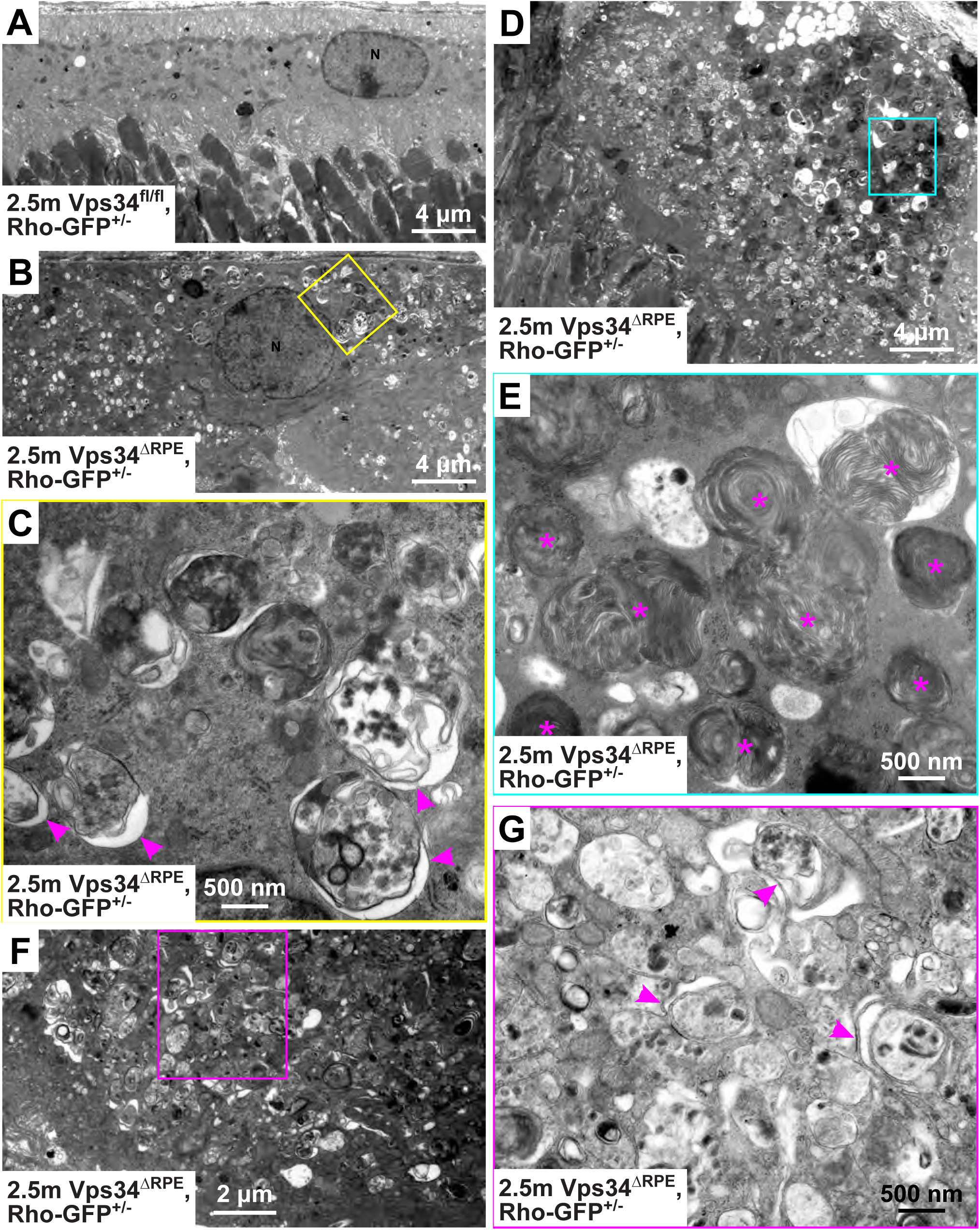
Phagosomes and autophagosomes accumulate in Vps34 KO RPE. TEM of 2.5-month-old control (Vps34^fl/fl^) (**A**) and KO (Vps34^ΔRPE^) (**B-G**) RPE. Rectangles indicate regions shown in higher-magnification images. Accumulation of various types of compartments was observed in different regions including POS-containing phagosomes (**E**, asterisks), and double-membrane and multilamellar autophagosomes (**C**, **G**, arrowheads).

**Supplemental Figure S3 (Related to Figure 5).**
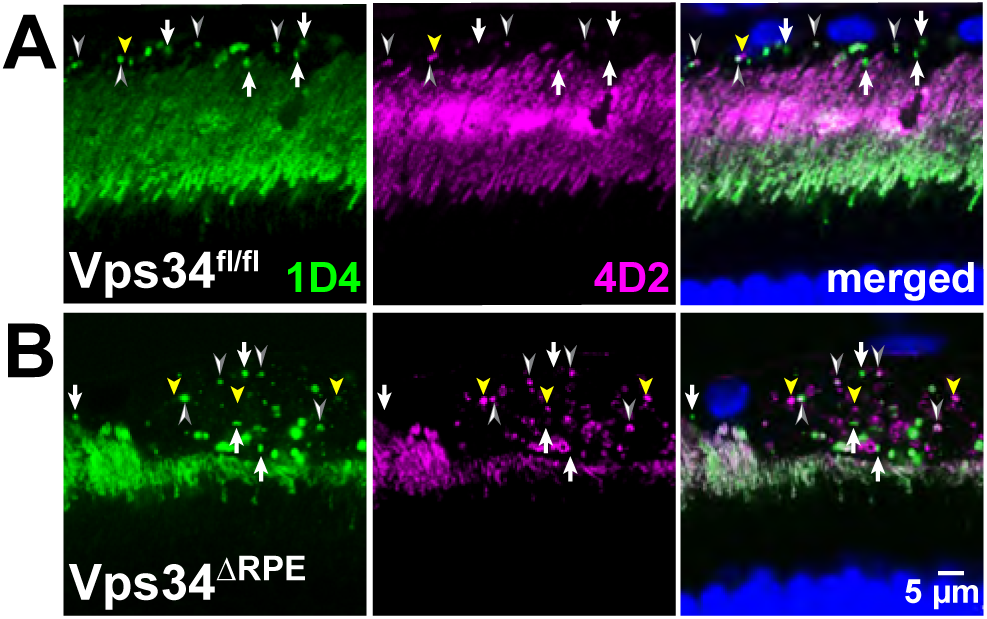
Degradation of Rho during phagosome maturation. Eyecup sections from control (**A**) or KO (**B**) animals were co-immunostained with Rho N-terminal antibody 4D2 and C-terminal antibody 1D4. In both genotypes, phagosomes had predominantly 1D4 staining (*white arrows*), clear co-staining with both antibodies (*thin, white arrowheads*) or predominantly 4D2 staining (*yellow arrowheads*).

**Figure S4 (Related to Figure 6).**
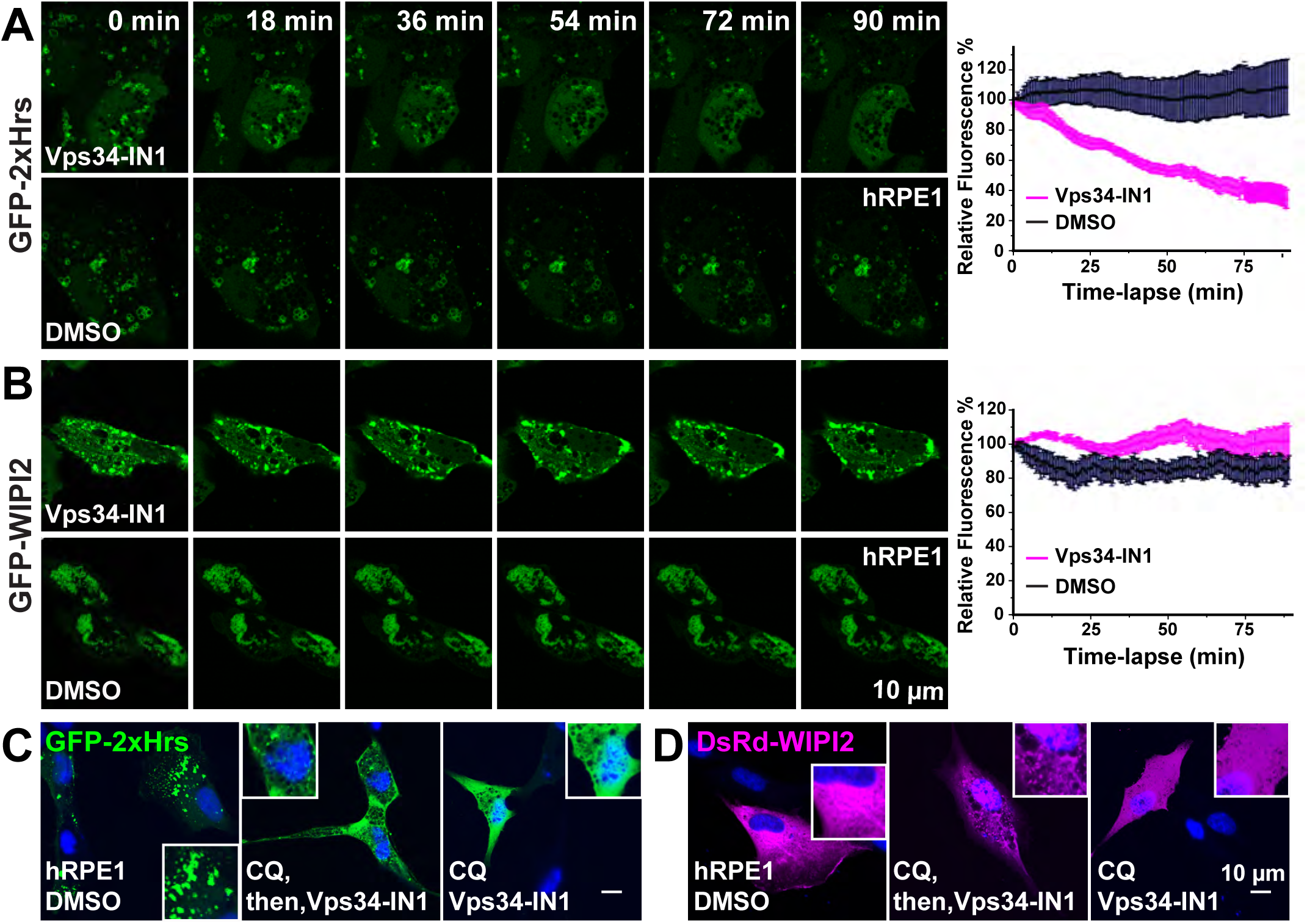
WIPI2 puncta formation requires PI(3)P, but localization is resistant to PI(3)P depletion. (**A**) hRPE1 cells were transfected with GFP-2xHrs or GFP-WIPI2 and treated with CQ, then either Vps34-IN1 or DMSO. Images of live cells were acquired at 30s intervals for 90m. (B) hRPE1 cells were transfected with GFP-2xHrs or dsRed-WIPI2, and treated with CQ overnight, followed by acute Vps34-IN1 exposure for 90 min, or with both CQ and Vps34-IN1 overnight. Following acute Vps34-IN1 treatment, 2xHrs puncta are disrupted, but WIPI2 puncta are not; following overnight Vps34-IN1 treatment, both probes are diffusely localized.

**Supplemental Figure S5 (Related to Figure 10).**
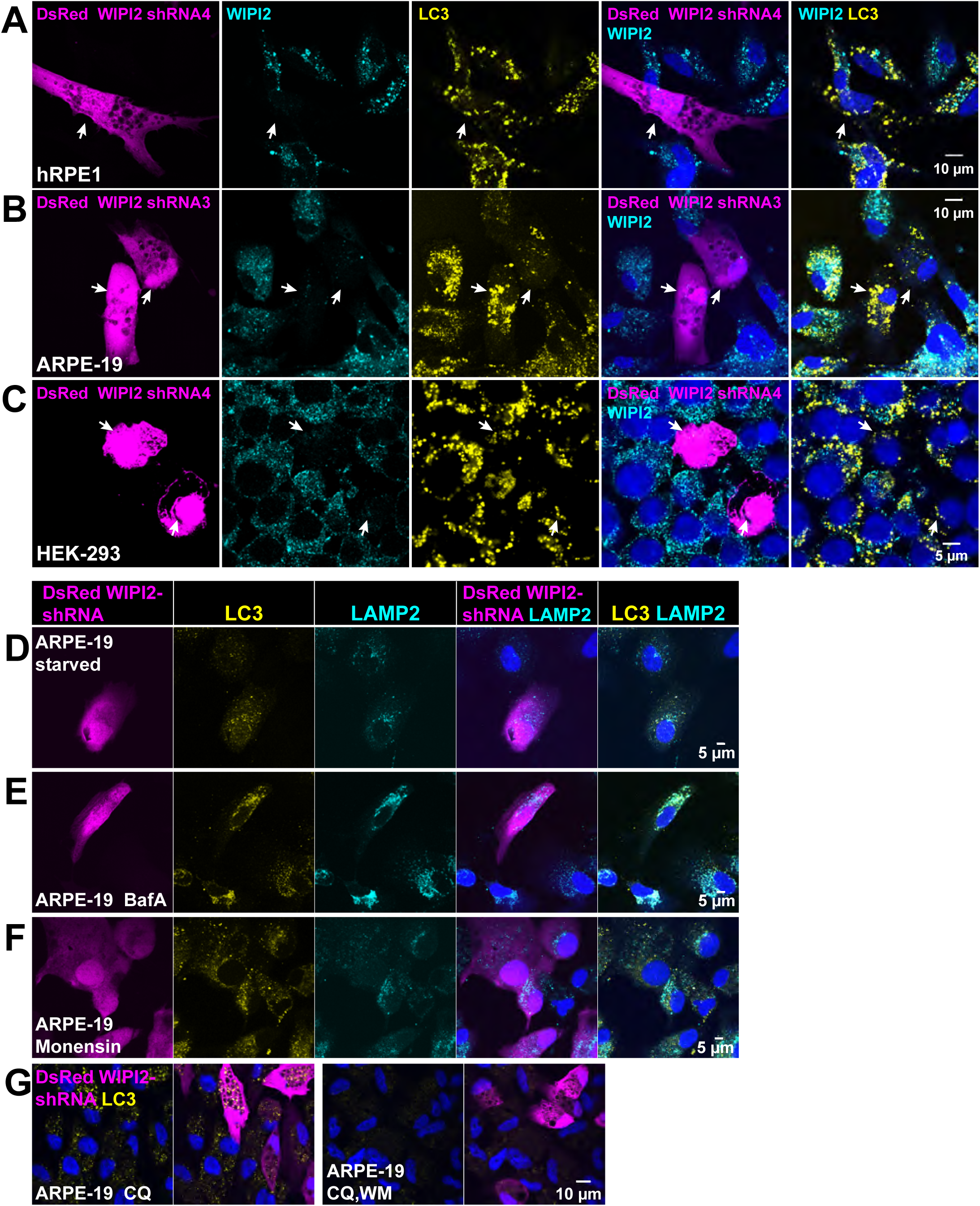
Effects of WIPI2 knockdown on LC3 puncta and on lysosome colocalization. hRPE1 (**A**) ARPE-19 (**B**) or HEK (**C**) cells were transfected with WIPI2 shRNA (marked by DsRed expression) and immunostained for WIPI2 and LC3. (**B**) Results showing LC3 puncta in WIPI2 knockdown ARPE-19 cells were reproduced with a second shRNA targeting a different sequence in WIPI2 than the one used in Fig. 10. **(D-G)** ARPE-19 cells were transfected with WIPI2 shRNA (marked by DsRed expression), and treated with starvation (**D**), bafilomycin A (**E**), or monensin (**F**), then immunostained for LC3 and LAMP2. LC3 puncta and LAMP1-positive lysosomes accumulate in each case, but lysosome co-localization with LC3 is much more pronounced in response to starvation or bafilomycin treatment than in response to monensin treatment. (**G**) ARPE-19 cells were transfected with WIPI2 shRNA, and treated overnight with CQ or CQ and wortmannin together.

